# Structural analysis of RIG-I-like receptors reveals ancient rules of engagement between diverse RNA helicases and TRIM ubiquitin ligases

**DOI:** 10.1101/2020.08.26.268649

**Authors:** Kazuki Kato, Sadeem Ahmad, Zixiang Zhu, Janet M. Young, Xin Mu, Sehoon Park, Harmit S. Malik, Sun Hur

## Abstract

RNA helicases and ubiquitin E3 ligases mediate many critical functions within cells, but their actions have been studied largely in distinct biological contexts. Here, we uncover evolutionarily conserved rules of engagement between RNA helicases and tripartite motif (TRIM) E3 ligases that lead to their functional coordination in vertebrate innate immunity. Using cryo-electron microscopy and biochemistry, we show that RIG-I-like receptors (RLRs), viral RNA receptors with helicase domains, interact with their cognate TRIM/TRIM-like E3 ligases through similar epitopes in the helicase domains. Their interactions are avidity-driven, restricting the actions of TRIM/TRIM-like proteins and consequent immune activation to RLR multimers. Mass-spectrometry and phylogeny-guided biochemical analyses further reveal that similar rules of engagement apply to diverse RNA helicases and TRIM/TRIM-like proteins. Our analyses thus reveal not only conserved substrates for TRIM proteins but also unexpectedly deep evolutionary connections between TRIM proteins and RNA helicases, thereby linking ubiquitin and RNA biology throughout animal evolution.

## Introduction

Innate immunity to invading pathogens involves tradeoffs between efficiency versus fidelity of responses. An inefficient response to an invading pathogen can be fatal, but an inappropriate immune reaction (*e.g*., in the absence of pathogen) can be equally harmful. A growing number of reports suggest that ubiquitin (Ub) modification of immune components provides a general means for fine-tuning the innate immune system (Zheng and Gao, 2019). Precise control of ubiquitination and consequent alteration of the target molecule’s stability or function allows another layer of regulation to achieve both fidelity and robustness in immune responses. Acting at the final step of the three-enzyme (E1, E2, and E3) Ub-transfer cascade, E3 ligases dictate ubiquitination target specificity. TRIM and TRIM-like proteins are an emerging class of E3 ligases that play important roles in innate immunity (Ozato et al., 2008; Versteeg et al., 2014). For substrate recognition, TRIM proteins often use carboxy-terminal Pry-Spry (Pspry) domains (Esposito et al., 2017), which can recognize distinct protein features from linear peptide sequences to three-dimensional protein structures (D’Cruz et al., 2013; Perfetto et al., 2013). Despite being present in nearly 100 human proteins, our fundamental understanding of how PSpry domains recognize substrates and enables TRIM proteins to regulate immune function is limited.

The recent discovery of TRIM-like protein RIPLET as the E3 ligase responsible for ubiquitination of RIG-I (Cadena et al., 2019b; Hayman et al., 2019; Oshiumi et al., 2013) revealed important insights into PSpry:substrate interactions. RIG-I is a conserved innate immune receptor that activates type I and III interferon (IFN) pathways upon binding to viral double-stranded RNA (dsRNA) (Hur, 2019; Yoneyama and Fujita, 2010). RIPLET is required to add K63-linked Ub chains (K63-Ub_n_) onto RIG-I (Cadena et al., 2019; Hayman et al., 2019; Oshiumi et al., 2013), which was previously shown to be a key trigger for RIG-I signal activation (Jiang et al., 2012; Zeng et al., 2010). K63-Ub_n_ promotes homo-tetramerization of RIG-I’s tandem CARD (2CARD) signaling domains, which in turn interact with the adaptor molecule MAVS to activate downstream signaling (Peisley et al., 2014; Wu et al., 2014). RIPLET selectively recognizes RIG-I in its filamentous oligomeric form, which assembles only upon its engagement with cognate dsRNA, and does not bind monomeric RIG-I free of RNA (Cadena et al., 2019). This filament-specific recognition, mediated by the bivalency of PSpry within a dimeric architecture of RIPLET, prevents constitutive ubiquitination and aberrant activation of RIG-I in the absence of viral infection (Cadena et al., 2019).

MDA5 is a RIG-I-like receptor (RLR) that performs distinct antiviral functions from RIG-I by detecting different groups of viral RNAs (Hur, 2019; Yoneyama and Fujita, 2010). Like RIG-I, MDA5 forms filaments upon binding to cognate viral dsRNAs and activates MAVS and the IFN-inducing pathway (Peisley et al., 2011; Wu et al., 2013). Moreover, K63-Ub_n_ also serves as a key trigger for MDA5 signaling (Jiang et al., 2012). Unlike RIG-I, however, ubiquitination of MDA5 is independent of RIPLET and instead requires the E3 ligase TRIM65 (Cadena et al., 2019; Kamanova et al., 2016; Lang et al., 2017). In this study, we found that TRIM65 recognizes MDA5 only in its filamentous state, revealing avidity-dependent RLR recognition as a common mechanism by which TRIM65 and RIPLET ensure tightly regulated antiviral signaling. We also found that both RIPLET and TRIM65 bind the same epitope in the helicase domains of the cognate RLRs, which nevertheless confers highly selective substrate recognition. Furthermore, our biochemical analyses showed that similar epitope in the helicase domain is utilized for avidity-dependent recognition by a broad range of TRIM proteins, revealing the conserved rules of engagement between TRIM proteins and helicases.

## Results

### TRIM65 selectively recognizes MDA5 filaments using PSpry bivalency

We first tested the importance of TRIM65 in MDA5 signaling (Kamanova et al., 2016; Lang et al., 2017). CRISPR-mediated knockdown (KD) of TRIM65 in 293T cells impaired MDA5 signaling upon stimulation with dsRNA mimetic, polyIC (Figure 1A). Conversely, TRIM65 complementation restored MDA5 signaling, as measured by IFNβ mRNA induction (Figure 1A). TRIM65 KD did not affect IFNβ induction upon stimulation with a RIG-I-specific RNA ligand (42 bp dsRNA with a 5’-triphosphate group), or upon overexpression of MAVS or STING (Figures S1A–C), suggesting that TRIM65 specifically acts on the MDA5 pathway. We also examined whether signaling activity of a gain-of-function mutant of MDA5 (G495R) relied on TRIM65. MDA5 G495R, which was identified from patients with the auto-inflammatory disease Aicardi-Goutières syndrome (Rice et al., 2014), is known to constitutively activate MDA5 signaling because it allows spontaneous filament formation of MDA5 on endogenous dsRNAs (Ahmad et al., 2018). We found that the signaling activity of G495R also depended on TRIM65 (Figure 1B). This result not only reinforces the requirement of TRIM65 for MDA5-dependent signaling, but also demonstrates that the TRIM65 requirement is independent of the source of the RNA ligand.

**Figure 1.**
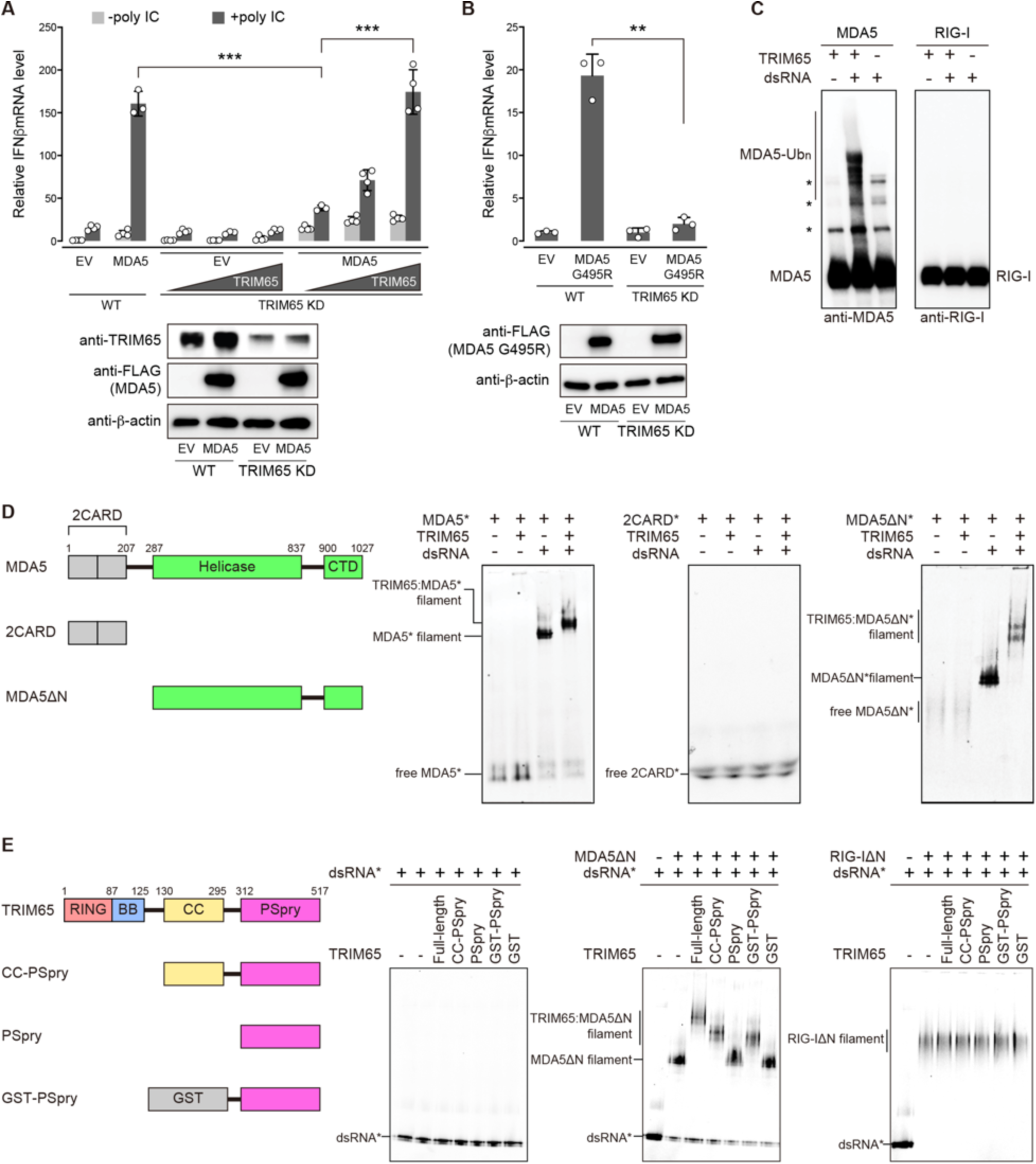
TRIM65 selectively recognizes MDA5 filaments using PSpry bivalency. (A) Relative signaling activity of MDA5 upon polyIC stimulation (0.5 μg), as measured by IFNβ mRNA induction. The signaling activity was compared in wild-type (WT) and TRIM65 KD 293T cells, with and without TRIM65 complementation. Since 293T expresses endogenous MDA5 poorly, cells were transfected with MDA5-expressing vectors. EV indicates empty vector. See Methods for experimental details. (B) Relative signaling activity of MDA5 G495R in WT and TRIM65 KD 293T cells. No exogenous RNA was introduced. (C) *In vitro* ubiquitination of MDA5 and RIG-I (500 nM) using TRIM65 (250 nM) in the presence and absence of dsRNA (2 ng/μl). 1012 bp and 112 bp dsRNAs were used for MDA5 and RIG-I, respectively, as the two receptors prefer different lengths of dsRNA for filament formation and immune stimulation. The Ubc13:Uev1a complex was used as E2. Reactions were analyzed by SDS-PAGE, followed by anti-MDA5 and anti-RIG-I immunoblots. (D) Native gel mobility shift assays to monitor MDA5:TRIM65 interaction. Fluorescein-labeled (*) full-length MDA5, 2CARD or MDA5ΔN (600 nM) were incubated with TRIM65 (300 nM) in the presence and absence of dsRNA (4 ng/μl). We used 112 bp dsRNA, instead of 1012 bp dsRNA, to clearly visualize the mobility shift upon TRIM65 binding. Fluorescein fluorescence was used for gel imaging. (E) Native gel mobility shift assays to monitor MDA5:TRIM65 interaction with TRIM65 domain mutants. Filaments of MDA5ΔN or RIG-IΔN (250 nM) were formed on Cy5-labeled (*) 112 bp dsRNA (1 ng/μl) and were incubated with TRIM65 or its truncation variants (300 nM). All data are representative of at least three independent experiments. Data in (a-b) are mean ± SD (n=3-4), and *P* values were calculated by two-tailed t-test (\*\*\**P*<0.001, \*\**P* <0.005).

Reconstitution of the ubiquitination system using purified proteins *in vitro* showed that TRIM65 directly ubiquitinates MDA5, but not RIG-I, under equivalent conditions (Figure 1C). We found that the Ub chains conjugated to MDA5 by E3 ligase TRIM65 and the E2 protein Ubc13:Uev1A were primarily K63-Ub_n_ (Figures S1D and S1E), which are required for MDA5 signal activation (Jiang et al., 2012). Mutation or deletion of TRIM65’s RING domain, which is required for E2 binding, impaired MDA5 ubiquitination (Figure S1F). Importantly, robust MDA5 ubiquitination by TRIM65 required dsRNA (Figure 1C), which mediates MDA5 filament formation and signal activation. Native gel mobility shift assays showed that TRIM65 binds MDA5 only in the presence of dsRNA (Figure 1D, left panel) whereas it does not bind RIG-I either in the presence or absence of dsRNA (Figure S1G). Since TRIM65 itself does not bind dsRNA (Figure 1E, left panel), our results suggest that the mobility shift of the MDA5 filament occurs via a direct interaction between the MDA5 filament and TRIM65.

Next, we investigated which domains mediate the TRIM65:MDA5 interaction. First, we tested the 2CARD domain of MDA5, and showed that it did not bind TRIM65, either in the presence or absence of dsRNA (Figure 1D, middle panel). However, we found that the 2CARD-deletion mutant (MDA5ΔN), which can form filaments on dsRNA, was able to bind TRIM65 in the presence of dsRNA (Figure 1D, right panel), implicating the helicase domain and/or CTD in the interaction. Similarly, neither the TRIM65 RING nor B-box domains were required for TRIM65 binding to MDA5 filaments, whereas both TRIM65 CC and PSpry domains were necessary (Figure 1E, middle panel; Figure S1H). However, we could replace CC with an artificial dimeric fusion protein, GST (Figure 1E, middle panel). Considering that TRIM65 CC is a constitutive dimer (Figure S1I), this result suggests that PSpry bivalency is both necessary and sufficient for MDA5 filament binding by TRIM65. Thus, not only is TRIM65 a highly specific E3 ligase for MDA5, just as RIPLET is for RIG-I (Cadena et al., 2019), but TRIM65 and RIPLET also share key aspects of substrate recognition, *i.e*. utilization of PSpry bivalency for selective recognition of RIG-I/MDA5 in the filamentous state.

### TRIM65 PSpry binds α1/α3 helices in the Hel2 domain of MDA5

We next investigated whether the functional similarity that we have uncovered between TRIM65:MDA5 and RIPLET:RIG-I interactions originates from a structural similarity in binding. We first determined cryo-EM structures of the MDA5 filaments on dsRNA bound by the PSpry domains from TRIM65. Since monomeric PSpry does not bind MDA5 filaments, we utilized an intramolecular fusion construct to increase the local concentration of PSpry and to promote its binding to MDA5. We fused TRIM65 PSpry (TRIM65^PSpry^) to the N- or C-terminus of MDA5ΔN through a flexible 38 amino acid-linker (Figure S2A). We confirmed that MDA5ΔN fused to TRIM65^PSpry^ can form filaments on dsRNA, independent of the fusion order (Figure S2B). In order to test whether the fused TRIM65^PSpry^ binds to MDA5ΔN in *cis*, we checked for the binding of the fusion constructs to GST-TRIM65^PSpry^ in *trans*. Our native gel mobility shift assay showed no binding of GST-TRIM65^PSpry^ to both N- and C-terminus fusion filaments, suggesting that the TRIM65 binding sites in these fusion constructs are pre-occupied by TRIM65^PSpry^ in *cis* (Figure S2C). For cryo-EM analysis, we chose the N-terminal fusion construct (TRIM65^PSpry^-MDA5ΔN) because of its superior protein stability compared to the C-terminal fusion construct.

Using fusion filaments formed on 1012 bp dsRNA (Figure 2A), we obtained cryo-EM data and computationally extracted trimeric segments of the filaments (see Supplementary Methods). From the resulting 15,306 trimeric segments, we reconstructed the cryo-EM map to 4.3 Å resolution (Figures 2B–C; Figures S2D–G and Table S1). We also determined the crystal structure of isolated TRIM65^PSpry^ to aid model building (1.9 Å, Table S2). The crystal structures of MDA5ΔN (Wu et al., 2013) and TRIM65^PSpry^ (this paper) could be fitted unambiguously in the reconstructed map (Figure 2B). Focused 3D classification and refinement of the central monomeric complex of MDA5ΔN:TRIM65^PSpry^ improved map quality for TRIM65^PSpry^ (4.3 Å, Figures S2E and S2F).

**Figure 2.**
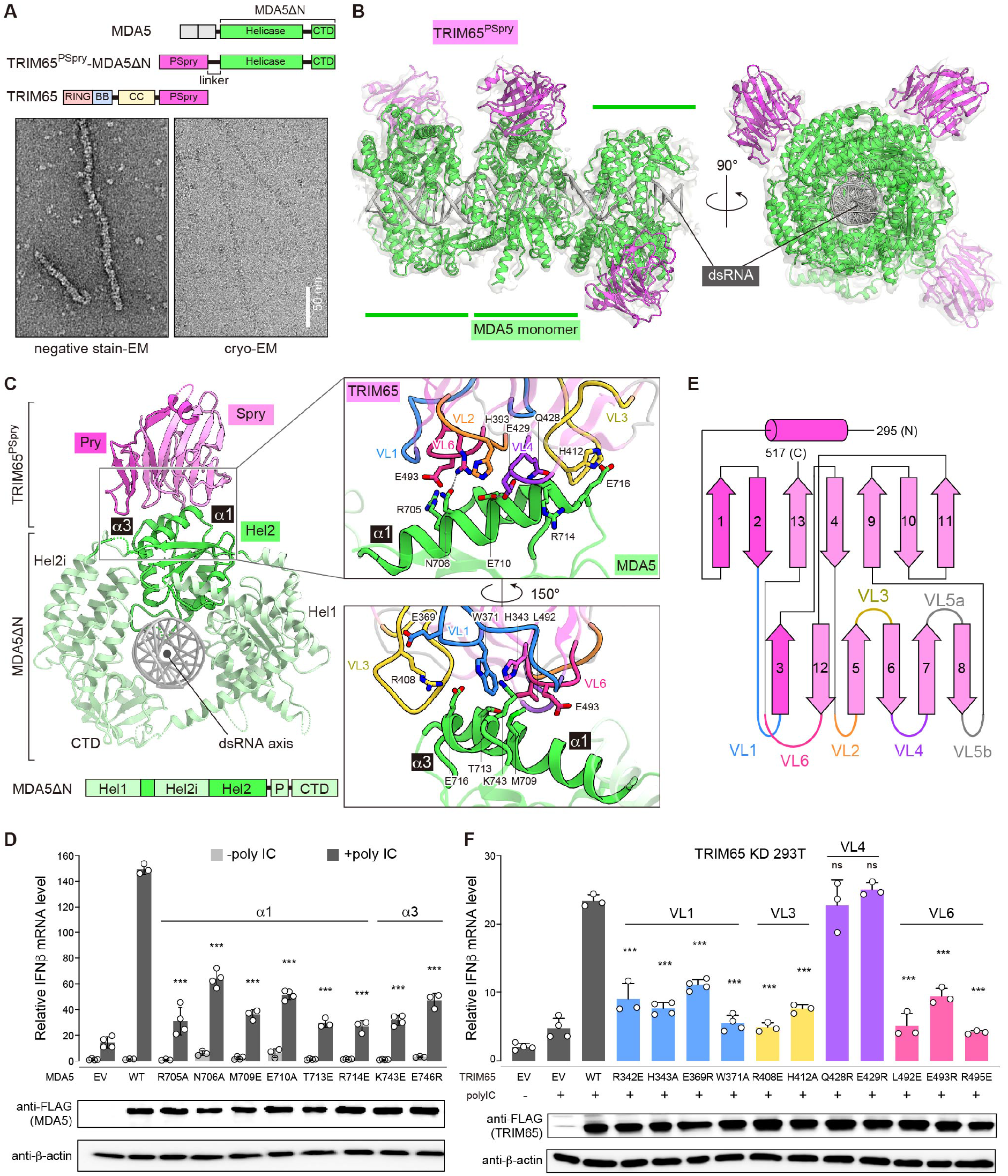
TRIM65 PSpry recognizes a1/a3 of MDA5 Hel2. (A) Top: Schematic of TRIM65^PSpry^-MDA5ΔN fusion construct. Bottom: Representative negative-stain (left) and cryo (right) electron micrographs of filaments formed by TRIM65^PSpry^-MDA5ΔN on 1012 bp dsRNA. (B) Cryo-EM map and ribbon model of the trimeric segment of the TRIM65^PSpry^-MDA5ΔN fusion filament. The cryo-EM map was low-pass filtered at 5 Å resolution and shown at 50% transparency. TRIM65^PSpry^, MDA5ΔN and dsRNA are colored magenta, green and grey, respectively. See also Figure S2 and Table S1 for details. (C) Top view of the central monomeric complex of MDA5ΔN:TRIM65^PSpry^. Subdomains of MDA5ΔN are colored according to the schematic below. Note that part of Hel2 is separated by the Hel2 insertion (Hel2i) domain. “P” indicates the pincer domain. Pry and Spry domains of TRIM65^PSpry^ are colored magenta and pink, respectively. Right, magnified views of the interface between TRIM65^PSpry^ and MDA5ΔN, where relevant variable loops (VLs) of TRIM65^PSpry^ are colored according to the secondary structure schematic in Figure 2E. Key interacting residues are shown in stick models. (D) Relative signaling activity of MDA5 with or without mutations in the TRIM65^PSpry^ binding interface as measured by IFNβ mRNA induction. 293T cells were transfected with MDA5-expressing vectors and stimulated with polyIC (0.5 μg). The amount of MDA5-expressing vector for each mutant was adjusted to ensure that the expression level is equivalent to WT. (E) Secondary structure representation of TRIM65^PSpry^ and definition of VLs. The Pry and Spry subdomains are distinguished by magenta and pink colors, respectively. (F) Relative signaling activity of WT MDA5 upon polyIC (0.5 μg) stimulation, as measured by IFNβ mRNA induction. TRIM65 KD 293T cells were transfected with plasmids expressing WT or mutant TRIM65. The amount of TRIM65-expressing vector for each mutant was adjusted to ensure that the expression level is equivalent to WT. All cells were transfected with equal amount of WT MDA5. Data in (D) and (F) are mean ± SD (n=3-4), and *P* values were calculated by comparing with the WT value using two-tailed t-test (\*\*\**P*<0.001; ns, not significant, *P*>0.1).

The structure showed that each TRIM65^PSpry^ interacts with an individual MDA5ΔN monomer with no contact across the filament interface (Figure 2B). TRIM65^PSpry^ binds two adjacent α1 (residues 700–715) and α3 (residues 743–747) helices within a subdomain (Hel2) of the MDA5 helicase domain (Figure 2C). Mutation analysis suggests that helices α1 and α3 are both important (Figure 2D). Structural comparison of Hel2 with (Wu et al., 2013) and without (Motz et al., 2013) dsRNA, and in the monomeric (Wu et al., 2013) vs. filamentous state (this study), suggests that neither Hel2 nor helices α1/α3 conformations change upon RNA binding or filament formation (Figure S3A). Consistent with these structural findings, forced dimerization of MDA5ΔN via GST fusion was sufficient to allow TRIM65 binding even in the absence of RNA, whereas cleavage of the GST tag impaired binding (Figure S3B). These data collectively suggest that filament specificity of TRIM65 is exclusively mediated by the bivalency requirement, instead of a direct readout of the filament structure.

TRIM65^PSpry^ displays a twisted β-sandwich fold (Figure 2C), which is common among the PSpry domains characterized so far (D’Cruz et al., 2013). It utilizes a cluster of variable loops (VLs) on one edge of the β-sandwich for MDA5 binding (Figures 2C and 2E). The MDA5 binding surface on TRIM65^PSpry^ is largely flat with a neutral electrostatic potential (Figure S3C), which may account for the low affinity interaction between individual TRIM65^PSpry^ and MDA5 molecules, and the requirement for avidity. The VLs from both Pry (VL1) and Spry subdomains (VL3, VL4 and VL6) contact MDA5^Hel2^ (Figure 2C, inset). By mutagenesis analysis, we confirmed that VL1, VL3 and VL6 in TRIM65^PSpry^ play important roles in MDA5 recognition, while residues in VL4 contacting MDA5 (e.g. Q428 and E429) may be less important (Figure 2F). Comparison with previous structures of two different PSpry domains in complex with their substrates (James et al., 2007; Woo et al., 2006) (Figure S3D) suggests that, while the PSpry domains commonly utilize VLs for substrate recognition, each PSpry employs a unique set of VLs with distinct sequences and lengths for binding diverse substrates.

### RIPLET recognizes RIG-I using a similar epitope in the helicase domain

We adopted a similar strategy to determine the cryo-EM structure of the human RIG-IΔN filament bound by RIPLET^PSpry^ (Figures 3A and 3B; Figure S4 and Table S1). The fusion filaments were formed on 112 bp dsRNA with a 5’-triphosphate group (Figure 3A), a preferred substrate for RIG-I. As with the TRIM65^PSpry^-MDA5ΔN filament, we first reconstructed a cryo-EM map for the trimeric segment of the RIPLET^PSpry^-RIG-IΔN filament (4.2 Å) and then performed focused local refinement of the central monomeric complex (3.9 Å). The maps were fitted with a previous crystal structure of human RIG-IΔN (Jiang et al., 2011) and a homology model of RIPLET^PSpry^ (Figure 3b).

**Figure 3.**
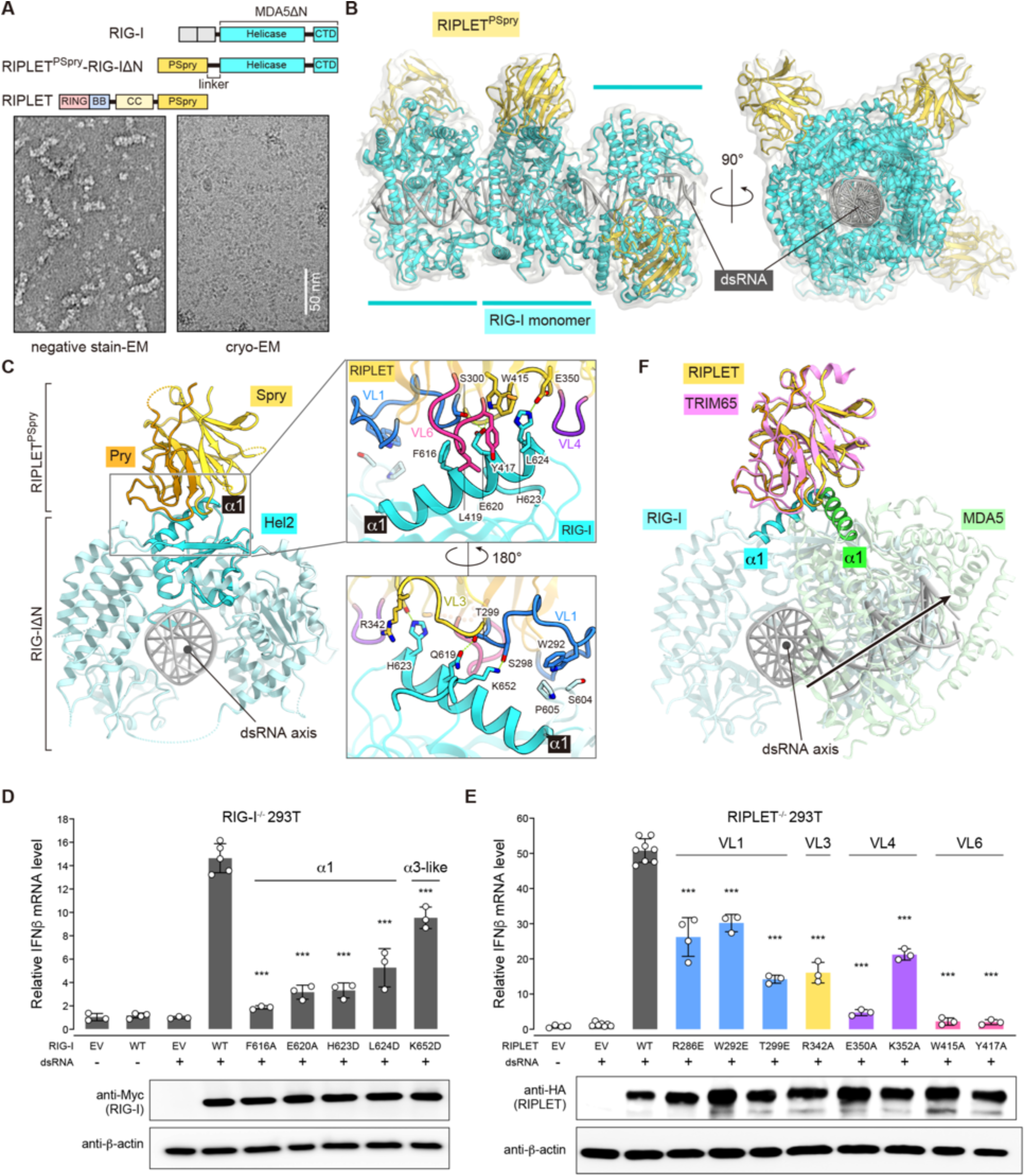
RIPLET PSpry recognizes RIG-I using a similar epitope as in the MDA5:TRIM65 complex, but the PSpry orientation differs. (A) Top: Schematic of RIPLET^PSpry^-RIG-IΔN fusion construct. Bottom: Representative negative-stain (left) and cryo (right) electron micrographs of filaments formed by RIPLET^PSpry^-RIG-IΔN on 112 bp dsRNA. (B) Cryo-EM map and ribbon model of the trimeric segment of the RIPLET^PSpry^-RIG-IΔN fusion filament. RIPLET^PSpry^, RIG-IΔN and dsRNA are colored yellow, cyan and grey, respectively. (C) Top view of the central monomeric complex of RIG-IΔN:RIPLET^PSpry^. The Hel2 subdomain and the rest of RIG-IΔN were colored cyan and light cyan, respectively. Pry and Spry domains of RIPLET^PSpry^ are colored orange and yellow, respectively. Right: magnified view of the interface between RIPLET^PSpry^ and RIG-IΔN, where relevant variable loops are displayed according to the color code in Figure 3E. Key interacting residues are shown in stick models. (D) Relative signaling activity of RIG-I with or without mutations in the RIPLET^PSpry^ binding interface as measured by IFNβ mRNA induction. RIG-I^-/-^ 293T cells (Shi et al., 2017) were transfected with RIG-I-expressing vectors and stimulated with 42 bp dsRNA with 5’-ppp (0.2 μg). The amount of RIG-I-expressing vector for each mutant was adjusted to ensure that the expression level is equivalent to WT. (E) Relative signaling activity of WT RIG-I upon stimulation with 5’-ppp-42 bp dsRNA (0.2 μg). RIPLET^7^’ 293T cells (Shi et al., 2017) were transfected with plasmids expressing WT or mutant RIPLET. The amount of RIPLET-expressing vector for each mutant was adjusted to ensure that the expression level is equivalent to WT. All cells were transfected with an equal amount of WT RIG-I. (F) Superposition of the MDA5ΔN:TRIM65^PSpry^ and RIG-IΔN:RIPLET^PSpry^ complexes by aligning PSpry domains of TRIM65 (pink) and RIPLET (yellow). With the exception of the a1 helices, MDA5 (green) and RIG-I (cyan) proteins are shown at 50% transparency. Data in (D) and (E) are mean ± SD (n=3-4), and *P* values were calculated by comparing with the WT value using two-tailed t-test (\*\*\**P*<0.001).

Intriguingly, the structure showed that RIPLET^PSpry^ also binds α1 (residues 609–624) and α3-like loop (residues 651–655) of RIG-I^Hel2^ (Figures 3C–3D) similar to TRIM65^PSpry^’s recognition of α1/α3 in MDA5^Hel2^. RIPLET^PSpry^ also utilizes VL1, VL3, VL4 and VL6 to contact RIG-I (Figures 3C and 3E). However, the relative orientation of RIPLET^PSpry^ differs from that of TRIM65^PSpry^ by ~60°, which is more evident when the two complex structures are superposed by aligning PSpry domains (Figure 3F). These divergent helicase-PSpry interaction modes are highly unexpected given the close relationship between RIG-I and MDA5 helicase domains, the overall similarity between RIPLET^PSpry^ and TRIM65^PSpry^ structures including the VL conformations (Figure 3F), and the functional parallelism in signaling between the two E3:RLR cognate pairs. Our structures uniquely highlight how PSpry domains can adapt to similar epitopes in divergent ways with only minimal perturbations in VLs.

### RLRs and their cognate TRIM proteins have co-evolutionary relationship

Sequence comparison among RIG-I and MDA5 homologs showed that the PSpry interface residues are relatively well-conserved within RIG-I orthologs and within MDA5 orthologs, but not between RIG-I and MDA5 (Figure 4A). This observation explains how RIPLET and TRIM65 achieve specificity towards RIG-I and MDA5, respectively, with little cross-reactivity. Intriguingly, some of the interface residues were not strictly conserved even among orthologs (Figure 4A; Figure S5). As a result, there is a limited cross-species reactivity between mouse and human proteins; mouse RIG-I and MDA5 are well-recognized by mouse RIPLET and TRIM65, respectively, but not as well by human RIPLET and TRIM65 (Figures 4B and 4C). This is despite the fact that mouse RIPLET and TRIM65 recognize both human and mouse RLRs equally well (Figures 4B and 4C). Analysis of the degree of conservation for the interface residues suggests that well-conserved residues in RIPLET/TRIM65 generally interact with well-conserved residues in RIG-I/MDA5, whereas less-conserved residues in RIPLET/TRIM65 interact with less-conserved residues in RIG-I/MDA5 (Figure 4D). These data collectively support our conclusion that interacting pairs of E3 ligases and RLRs have a co-evolutionary relationship.

**Figure 4.**
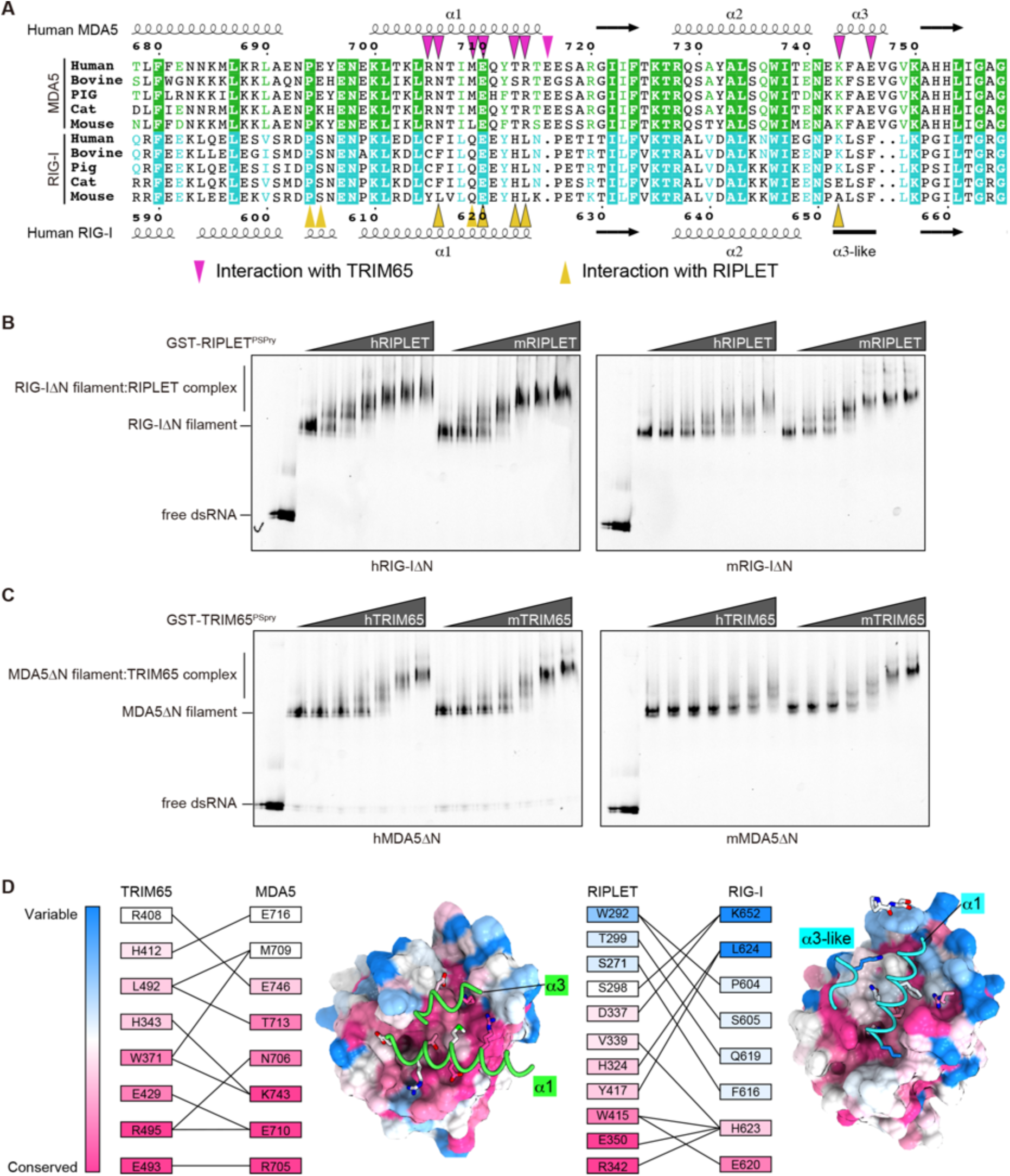
Conservation and co-evolution analysis of the RLR:PSpry interface. (A) Sequence alignment of orthologs of MDA5 and RIG-I near the TRIM65/RIPLET interface. Residues involved in the interaction with TRIM65 and RIPLET are indicated with triangles Sequences were aligned using Clustal Omega (http://www.ebi.ac.uk/Tools/msa/clustalo), and alignment figures were generated using ESPript3 (http://espript.ibcp.fr/ESPript/ESPript). (B) Native gel mobility shift assays of RIG-I filaments in the presence of RIPLET. RIG-I filaments were assembled by incubating Cy5-labeled 112 bp dsRNA (1 ng/μl) with human or mouse RIG-IΔN (hRIG-IΔN or mRIG-IΔN, 250 nM) in the presence of 2 mM ATP (Peisley et al., 2013). RIG-I filaments were then incubated with an increasing concentration (18-600 nM) of human or mouse RIPLET^PSpry^ (hRIPLET or mRIPLET) fused with GST, and the complex formation was analyzed by native PAGE using dsRNA fluorescence. (C) Native gel mobility shift assay of MDA5 filaments in the presence of TRIM65. Experiments were done as in (a) except that ATP was omitted in the reaction, because ATP promotes MDA5 filament disassembly (Peisley et al., 2011). (D) Degree of conservation for interacting residues in the MDA5:TRIM65 and RIG-I:RIPLET complexes. Conservation score was calculated using vertebrate protein sequences (see Methods) and Consurf web server (https://consurf.tau.ac.il/). The interacting residues are arranged in the descending order of conservation score (bottom to top), and the interaction pairs were indicated by connecting lines. Conservation score was also mapped onto the structures of PSpry using the program CueMol. The PSpry domains of TRIM65/RIPLET are shown in surface representation from the equivalent viewpoints, whereas a1/a3 helices of the bound RLRs are shown in cartoon representation.

### LGP2 is recognized by TRIM14 via PSpry bivalency and Hel2 epitope

We next turned our attention to LGP2, the third member of the RLR family with a helicase domain homologous to those of RIG-I and MDA5 (Sarkar et al., 2008) (Figure S6A). Like RIG-I and MDA5, LGP2 also forms filaments upon dsRNA binding (Uchikawa et al., 2016). However, unlike RIG-I and MDA5, LGP2 does not directly activate MAVS and the downstream pathway, but is thought to play a role in modulating RIG-I/MDA5 functions (Bruns and Horvath, 2015). Given the functional and structural similarity of TRIM65:MDA5 and RIPLET:RIG-I binding, we speculated that a related TRIM protein might bind and activate LGP2 in a similar manner. We used phylogenetic analysis to identify the human proteins most closely related to RIPLET and TRIM65 in their PSpry domains: TRIM47, TRIM25, TRIM16, TRIM16L, TRIM14 and BSPRY (Figure S6B). We were able to purify the PSpry domains of all these TRIM proteins except BSPRY as soluble GST fusions (Figure S6C). We tested these domains for their ability to interact with RIG-I, MDA5 and LGP2 filaments. Among the panel of PSpry domains tested, only TRIM65^PSpry^ and RIPLET^PSpry^ bound MDA5ΔN and RIG-IΔN filaments, respectively, again confirming the high selectivity of their engagement (Figure 5A). In contrast to an earlier report (Gack et al., 2007), but consistent with more recent findings (Cadena et al., 2019; Hayman et al., 2019), we found that TRIM25 does not interact with RIG-I (Figure 5A; Figure S6D).

**Figure 5.**
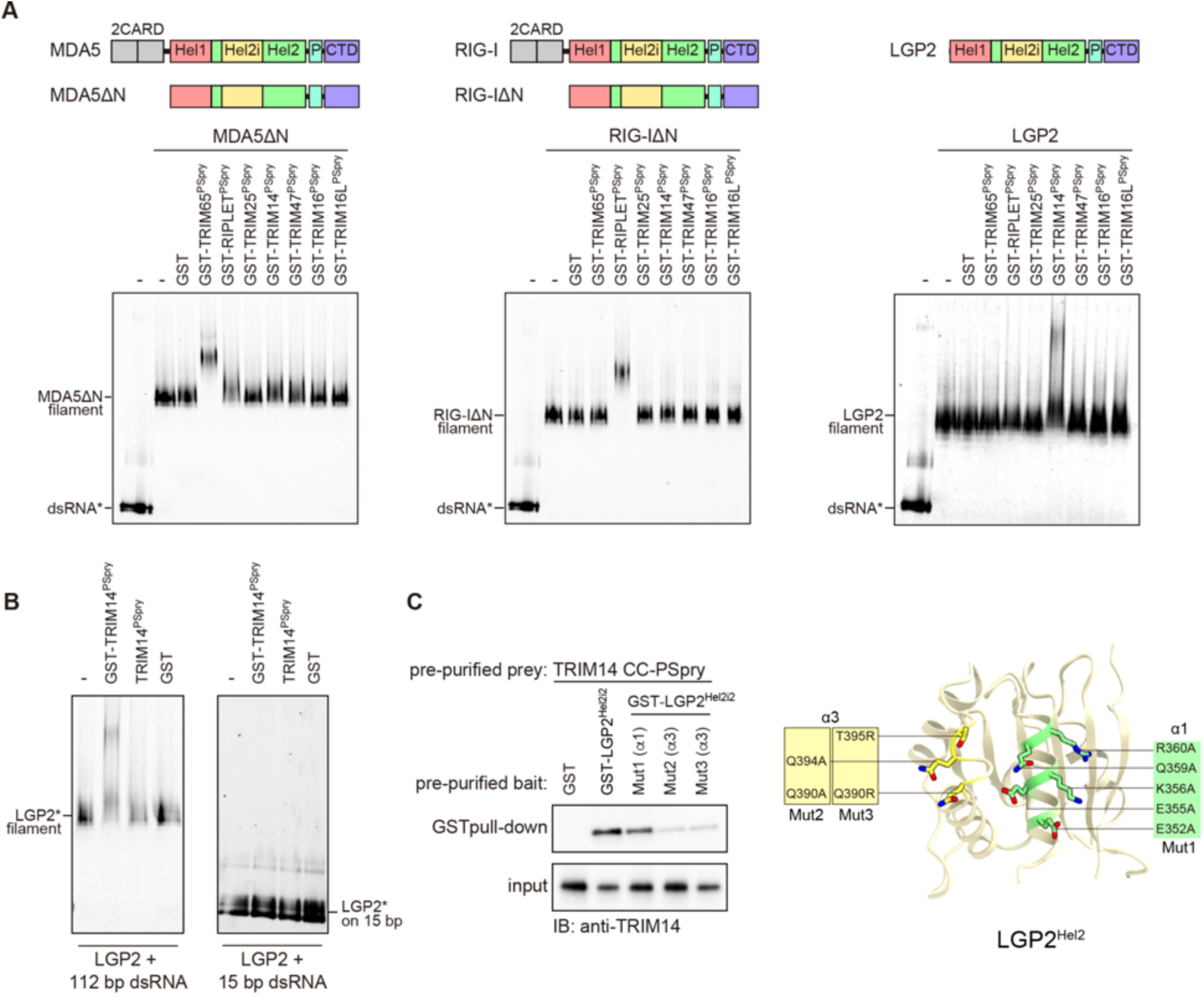
LGP2 is recognized by TRIM14 via PSpry bivalency and Hel2 epitope (A) Native gel mobility shift assay to test interactions between RLR filaments and a panel of TRIM^PSpry^ closely related to those of RIPLET and TRIM65 (see Figure S6). RLR filaments were formed by mixing MDA5ΔN, RIG-IΔN and LGP2 (250 nM) with Cy5-labeled (*) 112 bp dsRNA (1 ng/μl), and were incubated with TRIM^PSpry^ fused with GST (300 nM for RIG-I/MDA5 and 1.2 μM for LGP2). Cy5 fluorescence was used for gel imaging. (B) Mobility shift assay of LGP2 (0.6 μM) bound to 112 bp or 15 bp dsRNA (2.4 ng/μl) in the presence of GST-TRIM14^PSpry^, TRIM14^PSpry^ or GST alone (2.4 μM). (C) GST pull-down assay to examine the interaction between GST-LGP2^Hel2i2^ and TRIM14^CC-PSpry^. GST-LGP2^Hel2i2^ and TRIM14^CC-PSpry^ were recombinantly expressed in *E. coli* and purified prior to GST pull-down. The variants (Mut1-3) of LGP2^Hel2i2^ with mutations in α1 and α3 helices are defined on the right. Mutated residues are mapped onto the modeled human LGP2 Hel2, which was built using the Phyre2 server (http://www.sbg.bio.ic.ac.uk) with the crystal structure of chicken LGP2 (PDB:5JB2) as a template.

Using the same panel of GST-PSpry proteins, we also found that only TRIM14^PSpry^ is able to specifically bind LGP2 filaments (Figure 5A). TRIM14 was previously linked to the RIG-I pathway (Tan et al., 2017; Zhou et al., 2014). Given our finding that TRIM14 directly binds LGP2, but not RIG-IΔN, we deduce that the RIG-I-stimulatory function of TRIM14 may be indirectly mediated through its interaction with LGP2. TRIM14 binding to LGP2 also requires PSpry bivalency; TRIM14^PSpry^ without GST fusion does not bind LGP2 filaments (Figure 5B, left panel). Furthermore, GST-TRIM14^PSpry^ does not bind monomeric LGP2 on 15 bp dsRNA (Figure 5B, right panel). Given the evolutionary relatedness of LGP2 to RIG-I and MDA5, we next tested whether Hel2 of LGP2 is involved in TRIM14 binding. However, isolated LGP2 Hel2 was insoluble and required inclusion of the adjacent domain Hel2i to maintain solubility. Therefore, we purified LGP2 Hel2i-Hel2 (Hel2i2) fused to GST and performed GST pull-down. TRIM14^CC-PSpry^ co-purified with GST-LGP2^Hel2i2^, but not with GST alone (Figure 5C). Mutations in α3 helix, but not α1, of the Hel2 domain in LGP2^Hel2i2^ impaired the interaction between TRIM14^CC-PSpry^ and GST-LGP2^Hel2i2^ (Figure 5C). Overall, our results suggest that avidity-driven interaction of TRIM14 with LGP2 is mediated by an epitope similar to those of RIG-I and MDA5. Our findings highlight an ancient functional and structural similarity in TRIM-mediated binding of all three RLR proteins, which last had a common ancestor prior to the origin of bony vertebrates and the interferon-based immune system ~600 mya (Krause and Pestka, 2005; Sarkar et al., 2008).

### Distinct TRIM proteins recognize common epitopes in diverse helicases using PSpry bivalency

We next asked whether the helicase:PSpry interaction mode we identified with RLRs is more widespread among distantly related helicases. We investigated Dicer, which contains a helicase domain similar to those of RLRs (Kim et al., 2009) and, together with RLRs, belongs to the dsRNA-activated ATPases (DRAs) (Luo et al., 2013). Dicer is a conserved ribonuclease involved in RNA interference, which is considered one of the primary antiviral mechanisms that existed prior to the birth of the interferon system (Aguado et al., 2017; Maillard et al., 2019). To identify TRIM/TRIM-like proteins that interact with Dicer^Hel2^, we mixed purified GST-Dicer^Hel2i2^ with 293T lysate and subjected it to GST pull-down. We conjectured that GST fusion would mimic helicase dimerization and would allow identification of avidity-dependent interactors. Co-purified proteins were analyzed by mass-spectrometry and/or western blot analyses. Putative interaction partners containing PSpry domains were then recombinantly purified and re-analyzed for direct interactions with GST-Dicer^Hel2i2^. We found that TRIM25 copurifies with GST-Dicer^Hel2i2^ from 293T lysate, but not with GST (Figure 6A). The interaction is direct, as TRIM25^CC-PSpry^ and GST-Dicer^Hel2i2^ pre-purified from *E. coli* are also co-isolated by GST pull-down (Figure 6B). Neither TRIM25^PSpry^ nor TRIM25^CC^ alone can bind GST-Dicer^Hel2i2^ (Figure 6B), consistent with the importance of PSpry bivalency for their interactions. To examine whether the Hel2 regions equivalent to α1 and α3 of RLRs are involved in the interaction with TRIM25^CC-PSpry^, we made mutations in and around α1 helix and α3-like loop, which include linker residues (designated L1 and L2, Figure S6E) that are uniquely present in Dicer but not RLRs. Although the importance of α3-like loop could not be tested due to insolubility of the mutants, L1 adjacent to a3-like loop and L2 adjacent to a1 helix were both found important for TRIM25 binding (Figure 6B). These results suggest that TRIM25 binds Dicer by recognizing a similar region of Hel2 as in RLRs, again using PSpry bivalency. Thus, the mode of TRIM-helicase interaction that we have uncovered significantly preceded the origin of the interferon-based immune system (Krause and Pestka, 2005).

**Figure 6.**
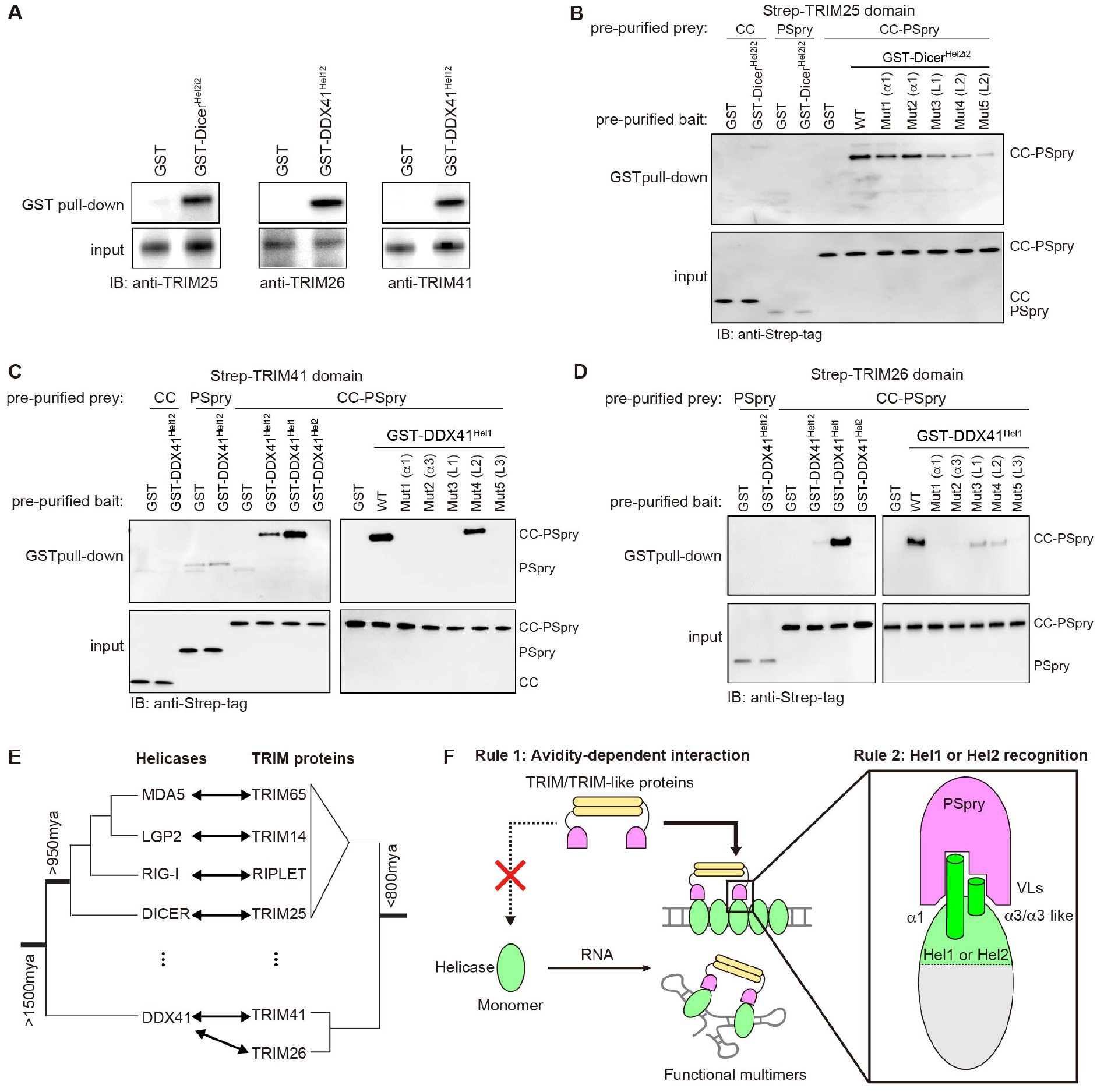
Multiple TRIM proteins recognize common epitopes in helicases beyond RLRs. (A) GST pull-down assays to identify interacting TRIM partners of Dicer and DDX41 helicase domains. GST-fused Hel2i-Hel2 of Dicer (GST-Dicer^Hel2i2^) and Hel1-Hel2 of DDX41 (GST-DDX41^Hel12^) were recombinantly purified, and mixed with 293T lysate for GST pulldown. (B) GST pull-down assay to examine direct interaction between GST-Dicer^Hel2i2^ and TRIM25. GST-Dicer^Hel2i2^ and TRIM25 (CC, PSpry or CC-PSpry) were recombinantly expressed in *E. coli* and purified prior to GST pull-down. The variants (Mut1-5) of Dicer^Hel2i2^ with mutations in a1 or L1 and L2 are defined in Figure S6E. (C-D) GST pull-down assays to examine direct interaction between GST-DDX41^Hel12^ and TRIM41^CC-PSpry^ (C) or TRIM26^CC-PSpry^ (D). GST-DDX41 (Hel12, Hel1 or Hel2) and TRIM41 or TRIM26 (CC, PSpry or CC-PSpry) were recombinantly expressed in *E. coli* and purified prior to GST pull-down. TRIM26 CC was insoluble and thus was not included in GST pull-down. The mutations (Mut1-5) in DDX41^Hel1^ are defined in Figure S6E. (E) Schematic phylogenetic trees of helicases and TRIM/TRIM-like proteins discussed in this study. See Figures S6A and S6B for full trees. (F) Cartoon summarizing the two rules of engagement between helicases and TRIM proteins. Rule 1: PSpry bivalency allows TRIMs to specifically recognize the cognate helicases in the multimeric form vs. monomeric form. This restricts the actions of TRIMs to the helicases in complex with certain RNAs that can induce the helicase multimerization. Multimerization can be in the form of filaments (as for RLRs) or other structures shaped by the conformation of the bound RNAs. Rule 2: Individual PSpry recognizes regions in or near α1/α3 of Hel1 or Hel2. Utilization of the common epitope by a wide range of helicases for their interaction with PSpry suggests an ancient evolutionary relationship between helicase and PSpry domains.

Finally, we investigated an even more divergent helicase, DDX41. Although DDX41 belongs to the same superfamily of helicases as Dicer and RLRs (Singleton et al., 2007), it is a DEAD-box helicase rather than a DRA (Luo et al., 2013), and diverged from DRAs more than 1500 mya (Figure 6E). DDX41 was previously known to be involved in splicing, translation and many other cellular RNA processes (Jiang et al., 2017; Peters et al., 2017; Polprasert et al., 2015), but was recently shown to participate in foreign DNA sensing, albeit through a poorly understood mechanism (Parvatiyar et al., 2012; Quynh et al., 2015; Zhang et al., 2011). DDX41 does not contain a Hel2i domain, but harbors Hel1 and Hel2 that share the RecA-like fold (Omura et al., 2016). Given the structural similarity between Hel1 and Hel2, we expanded our analysis to search for partners of DDX41 Hel1 as well as Hel2. We purified DDX41 Hel1-Hel2 fused with GST (GST-DDX41^Hel12^) and subjected it to the same pull-down and mass-spectrometry analysis as GST-Dicer^Hel2i2^. We found that two TRIM proteins, TRIM26 and TRIM41, specifically copurified with GST-DDX41^Hel12^ from 293T lysate (Figure 6A). Using pre-purified proteins, we showed that these interactions are direct, dependent on PSpry bivalency, and mediated by regions in DDX41 Hel1 equivalent to α1/α3 of Hel2 (Figures 6C and 6D). Our findings thus suggest that the rules of PSpry engagement we identified with RLRs and Dicer are not restricted to DRAs, but are shared with other helicases that diverged close to the origin of eukaryotes, nearly a billion years ago (Figure 6E).

## Discussion

We discovered two molecular principles that govern interactions between TRIM/TRIM-like proteins and RNA helicases (Figure 6F). The first principle is avidity-dependent interaction; many TRIM/TRIM-like proteins engage with the cognate helicases only in a multimeric state. The importance of this principle is best illustrated with TRIM65 and RIPLET, which respectively recognize filamentous MDA5 and RIG-I in the presence of foreign dsRNAs, but not in the monomeric, resting state. This enables tight control of MDA5/RIG-I-mediated immune signaling and ensures activation only upon viral infection. This avidity-dependent interaction also applies to other helicases (LGP2, Dicer and DDX41). We identified TRIM partners for these helicases and showed that their interactions are also dependent on PSpry bivalency. LGP2 forms filamentous oligomers along dsRNA, like MDA5 and RIG-I. Exactly how and when Dicer and DDX41 form multimers is yet unclear. Given the similarity between the helicase domains of Dicer and RLRs, Dicer may form filamentous oligomers on long dsRNA. Alternatively, multiple Dicer and DDX41 molecules may be indirectly bridged by certain RNAs, allowing bivalent binding of cognate TRIM proteins (Figure 6F). Regardless of the mechanism for helicase multimerization, our findings reveal a general rule by which TRIM/TRIM-like proteins regulate their engagement with helicases. They also raise an intriguing model that TRIM/TRIM-like proteins may function as a new type of reader for RNA structure and conformation by monitoring helicase multimerization.

The second principle we have uncovered is that regions of Hel1 or Hel2 near a1/a3 are commonly recognized by a broad range of TRIM/TRIM-like proteins harboring PSpry domains (Figure 6F). Our cryo-EM structures of the TRIM65:MDA5 and RIPLET:RIG-I complexes revealed that TRIM65 and RIPLET PSpry domains both recognize a1/a3 of cognate helicases, albeit adopting two distinct orientations. Remarkably, our analysis of LGP2, Dicer and DDX41 and their TRIM partners suggests that these interactions also use similar epitopes in the helicase domains. These observations raise questions about the evolutionary origins of this widespread recognition of common epitopes in Hel1/Hel2 domains of helicases by TRIM proteins. Phylogenetic analysis suggests that DDX41, Dicer and the ancestral RLR helicases diverged from one another significantly earlier than did their partner TRIMs (Figure 6E), so helicase divergence could not have driven concurrent TRIM evolution. It is possible that multiple TRIMs independently evolved to recognize the Hel1/Hel2 epitope region, perhaps because it allows sufficient specificity and accessibility when the helicase domain multimerizes. Given the low likelihood that convergent evolution would arrive at such similar binding determinants, we favor the alternate hypothesis that recognition of Hel1 or Hel2 epitopes might have been a property of an ancestral PSpry-containing TRIM protein. This ability may have been preserved in some descendant extant TRIM/TRIM-like proteins, whereas other lineages may have diverged to recognize non-helicase substrates. Regardless of whether the multiple TRIM-helicase interactions we have uncovered have resulted from convergent or divergent evolution, the recurrence of this interaction in multiple branches of TRIMs and helicases clearly suggests a widespread existence of such interactions and their co-option by different helicases to acquire Ub-mediated regulatory functions. Given that the functions and mechanisms of many helicases and TRIM proteins remain poorly understood, our findings provide a new mechanistic link between the two protein families in innate immunity and beyond.

## Acknowledgement

We acknowledge Dr. Adam Frost (UCSF) for the guidance with cryo-EM analysis. KK was supported by the Uehara Research Fellowship and a Cancer Research Institute Postdoctoral Fellowship. ZZ was supported by the China Scholarship Council (no. 2018032500003). JMY and HSM are supported by grants from the Mathers Foundation and NIAID (HARC [HIV Accessory and Regulatory Complexes] center, P50 AI082250, PI Nevan Krogan, subaward to HSM) and an HHMI Investigator award. SH acknowledges NIH grants (R01AI154653, R01AI111784 and DP1AI152074). The cryo-EM grids were screened at PNCC at OHSU (U24GM129547), and cryo-EM data were collected at NCI-NCEF at the Frederick National Laboratory for Cancer Research (under contract HSSN261200800001E) and The Harvard Cryo-EM Center for Structural Biology at Harvard Medical School.

---

**Figure S1.**
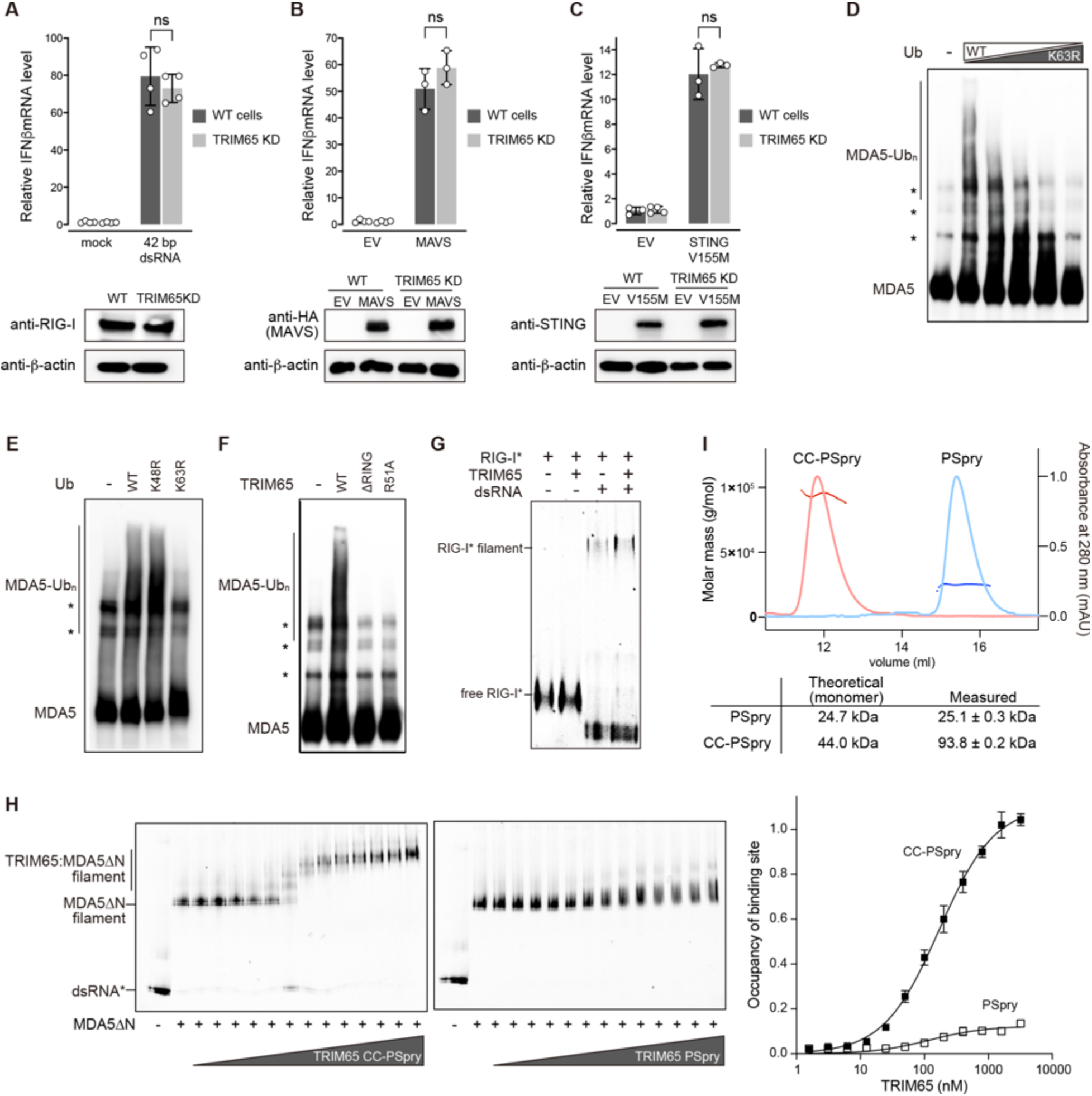
TRIM65 recognizes filamentous MDA5 and conjugates K63-Ub_n_. (A-C) Relative signaling activity of RIG-I (A), MAVS (B) and STING (B) in WT and TRIM65 KD 293T cells. Endogenous RIG-I was stimulated with 42 bp dsRNA with a 5’-triphosphate group (5’-ppp, 50 ng), which is known to specifically activate RIG-I (Cadena et al., 2019b). MAVS and STING pathways were activated by ectopic expression of wild-type MAVS (25 ng) and a constitutively active variant (V155M) of STING (0.75 μg) (Jeremiah et al., 2014). Data are mean ± SD (n=3-4), and *P* values were calculated by two-tailed t-test (ns, not significant; *P*>0.1). (D) *In vitro* ubiquitination assay of MDA5 with an increasing ratio of K63R to wild-type Ub. Total concentration of Ub was kept constant at 20 μM. All reactions contained 1012 bp dsRNA (2 ng/μl). Reactions were performed as in Figure 1C and were analyzed by anti-MDA5 western blot. (E) *In vitro* ubiquitination assay of MDA5 with WT, K48R or K63R Ub (20 μM). (F) *In vitro* ubiquitination assay of MDA5 using wild-type TRIM65, ΔRING or R51A. R51 in TRIM65 corresponds to a conserved E2 binding residue in RING domains (Metzger et al., 2014). (G) Native gel mobility shift assays to test RIG-I:TRIM65 interaction. Fluorescein-labeled (*) full-length RIG-I (600 nM) was incubated with TRIM65 (300 nM) in the presence and absence of 112 bp dsRNA (4 ng/μl). Fluorescein fluorescence was used for gel imaging. The result shows no binding between RIG-I and TRIM65, with or without dsRNA. (H) Comparison between TRIM65 CC-PSpry and PSpry for MDA5 filament binding. MDA5ΔN filament was formed by mixing Cy5-labeled (*) 112 bp dsRNA (1 ng/μl) and MDA5ΔN (250 nM). The filament was then incubated with an increasing concentration of CC-PSpry or PSpry (1.57-6400 nM), and the complex formation was analyzed by native PAGE. Right: quantitative analysis of the mobility shift assay (mean ± SD, n=3), which showed that TRIM65 CC-PSpry has *K*_D_ of 170.6 nM. *K*_D_ could not be determined for PSpry due to inefficient mobility shift at all concentrations tested. (I) Size-exclusion chromatography (SEC) coupled multi-angle light scattering (MALS) analysis of TRIM65 CC-PSpry and PSpry. Measured and theoretical molecular weights of the proteins are indicated below.

**Figure S2.**
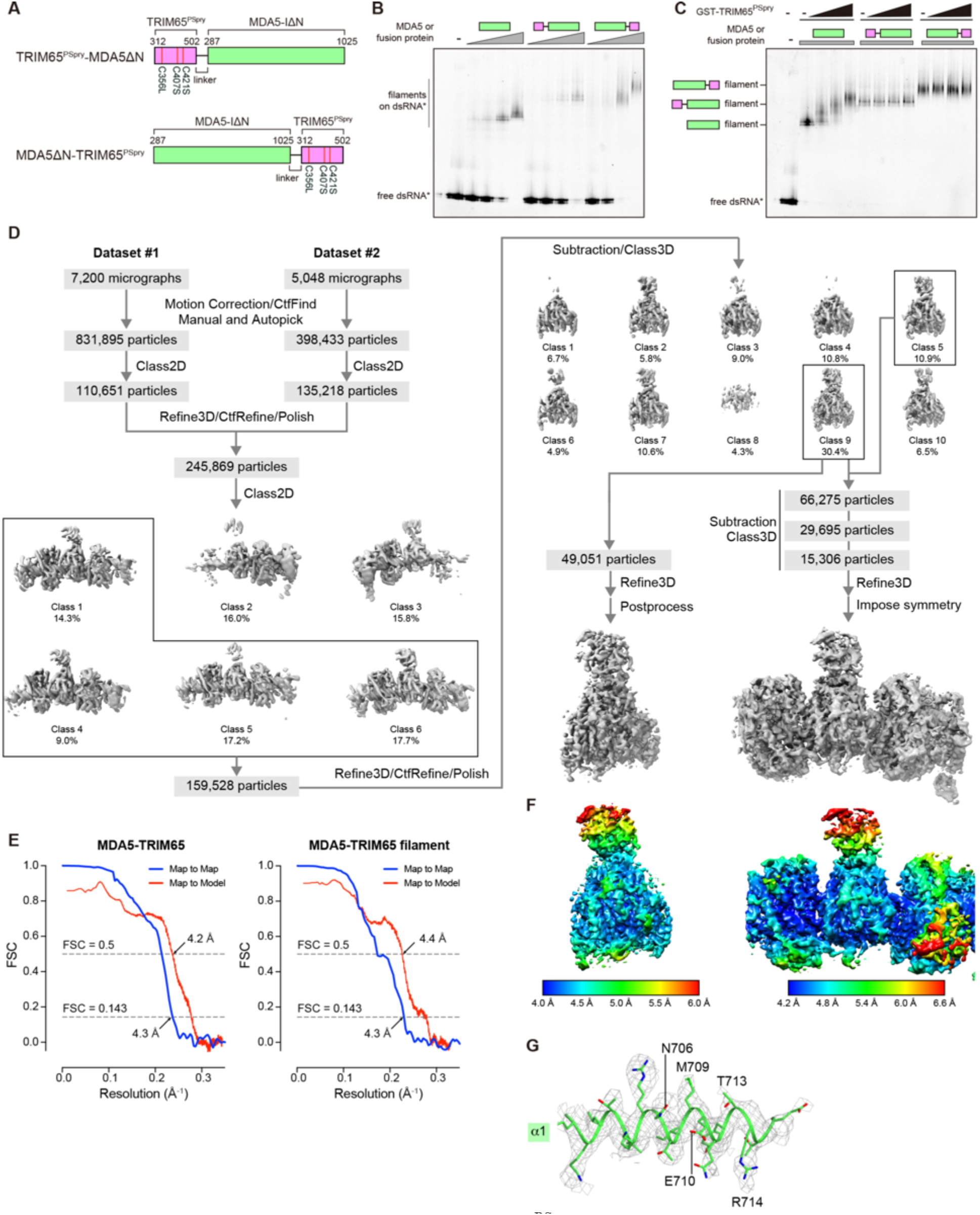
Cryo-EM data processing for the TRIM65^PSpry^:MDA5ΔN complex. (A) Schematic of TRIM65^PSpry^ fused with MDA5ΔN at the N- or C-terminus. The 38 amino acid linker was derived from the multi-cloning site of pET50b vector. Three Cys (C356, C407 and C421) in TRIM65^PSpry^ were mutated to improve the protein stability. (B) Filament formation of MDA5ΔN, TRIM65^PSpry^-MDA5ΔN and MDA5ΔN-TRIM65^PSpry^ on 112 bp dsRNA. The mobility shift of Cy5-labeled (*) dsRNA (1 ng/μl) was monitored by native PAGE with an increasing concentration (62.5, 125, 250 and 500 nM) of MDA5ΔN or the fusion proteins. Cy5 fluorescence was used for gel imaging. (C) GST-PSpry binding assay. Filaments of MDA5ΔN, TRIM65^PSpry^-MDA5ΔN and MDA5ΔN-TRIM65^PSpry^ were formed on Cy5-labeled (*) 112 bp dsRNA as in (b) and their mobilities were monitored by PAGE with an increasing concentration (72.5, 150 and 300 nM) of GST-TRIM65^PSpry^. (D) Cryo-EM image processing workflow (see details in Supplementary Methods). (E) Fourier shell correlation (FSC) curve for the monomeric (left panel) and filamentous (right panel) MDA5:TRIM65 complex. Map-to-Map FSC curve was calculated between the two independently refined half-maps after masking (blue line), and the overall resolution was determined by gold standard FSC=0.143 criterion. Map-to-Model FSC was calculated between the refined atomic models and maps (red line). (F) Local resolution for the maps of the monomeric (left panel) and filamentous (right panel) MDA5:TRIM65 complex. Local resolution was calculated by Relion, and resolution range was indicated according to the color bar. (G) Cryo-EM density map for the a1 helix of MDA5.

**Figure S3.**
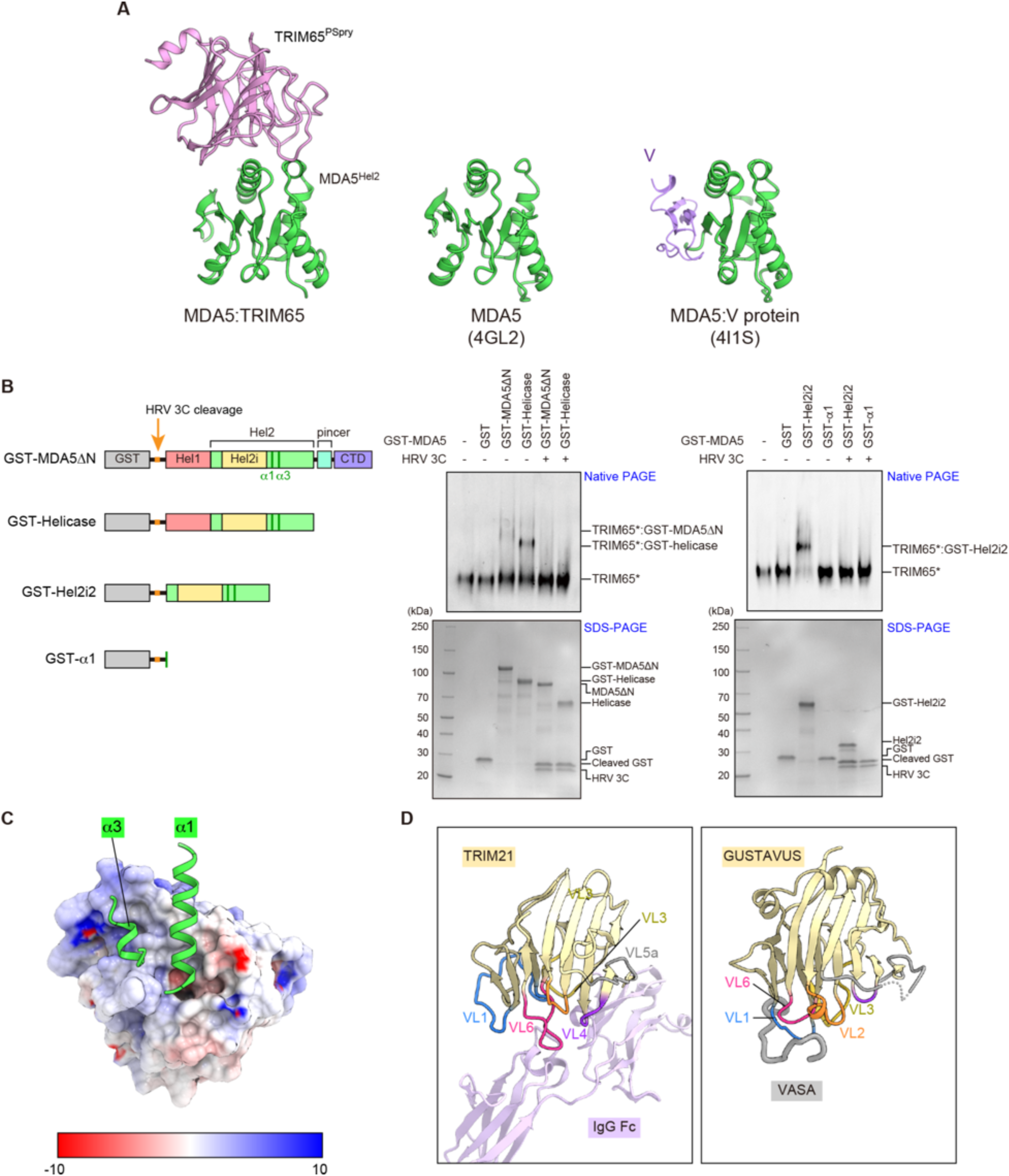
Analysis of the MDA5:TRIM65 complex structure. (A) Comparison of the MDA5 Hel2 structures in the filamentous state with dsRNA and TRIM65 (left, this study), in the monomeric state with dsRNA (center, PDB: 4GL2) and in complex with the viral protein V without dsRNA (right, PDB: 4I1S). The Cα RMSD between the left and center Hel2 structures, and between left and right Hel2 structures are 1.54 Å and 1.09Å, respectively. (B) Native gel shift assays to examine the interaction between MDA5 and TRIM65 in the absence of dsRNA. TRIM65 was N-terminally labeled with fluorescein (*) for visualization in the native gel. Mobility shift of TRIM65 (CC-PSpry, 0.8 μM) was monitored upon incubation with MDA5 fused with GST (GST-MDA5, 1.6 μM). Various MDA5 constructs were used to confirm the interaction between MDA5^Hel2^ and TRIM65: MDA5ΔN, helicase domain, Hel2i-Hel2 (Hel2i2) and isolated a1 helix. Note that isolated Hel2 could not be tested due to its insolubility. We instead used Hel2i2, which was soluble. The GST tag is cleavable by the HRV 3C protease, allowing comparison of the MDA5:TRIM65 interaction in the monomeric (without GST) vs. dimeric (with GST) states. The results showed that MDA5ΔN, helicase domain and Hel2i2 all bind TRIM65 in a manner dependent on GST fusion. Isolated a1 helix did not bind TRIM65, with or without GST, suggesting that a3 is also required. All proteins were recombinantly expressed in *E. coli* and purified to homogeneity. Bottom: Input samples analyzed by SDS-PAGE and Coommassie Brilliant Blue (CBB) stain. (C) Electrostatic potential of TRIM65^PSpry^ in surface representation. The a1/a3 helices of MDA5ΔN bound by TRIM65^PSpry^ are shown in ribbon representation (green). (D) Structures of other PSpry domains in complex with their respective substrates. Left: TRIM21^PSpry^ in complex with IgG Fc (PDB:2IWG (James et al., 2007)). Right: GUSTAVUS^PSpry^ in complex with a peptide isolated from VASA (PDB:2IHS (Woo et al., 2006)). VLs involved in substrate recognition are indicated by VL labels. TRIM21^PSpry^ utilizes VL1 and 3-6 to recognize a domain-domain interface of the IgG antibody, whereas GUSTAVUS^PSpry^ utilizes VL1-3 and 6 to recognize a linear peptide in VASA. Note that VASA is a helicase, but the PSpry epitope resides outside the helicase domain.

**Figure S4.**
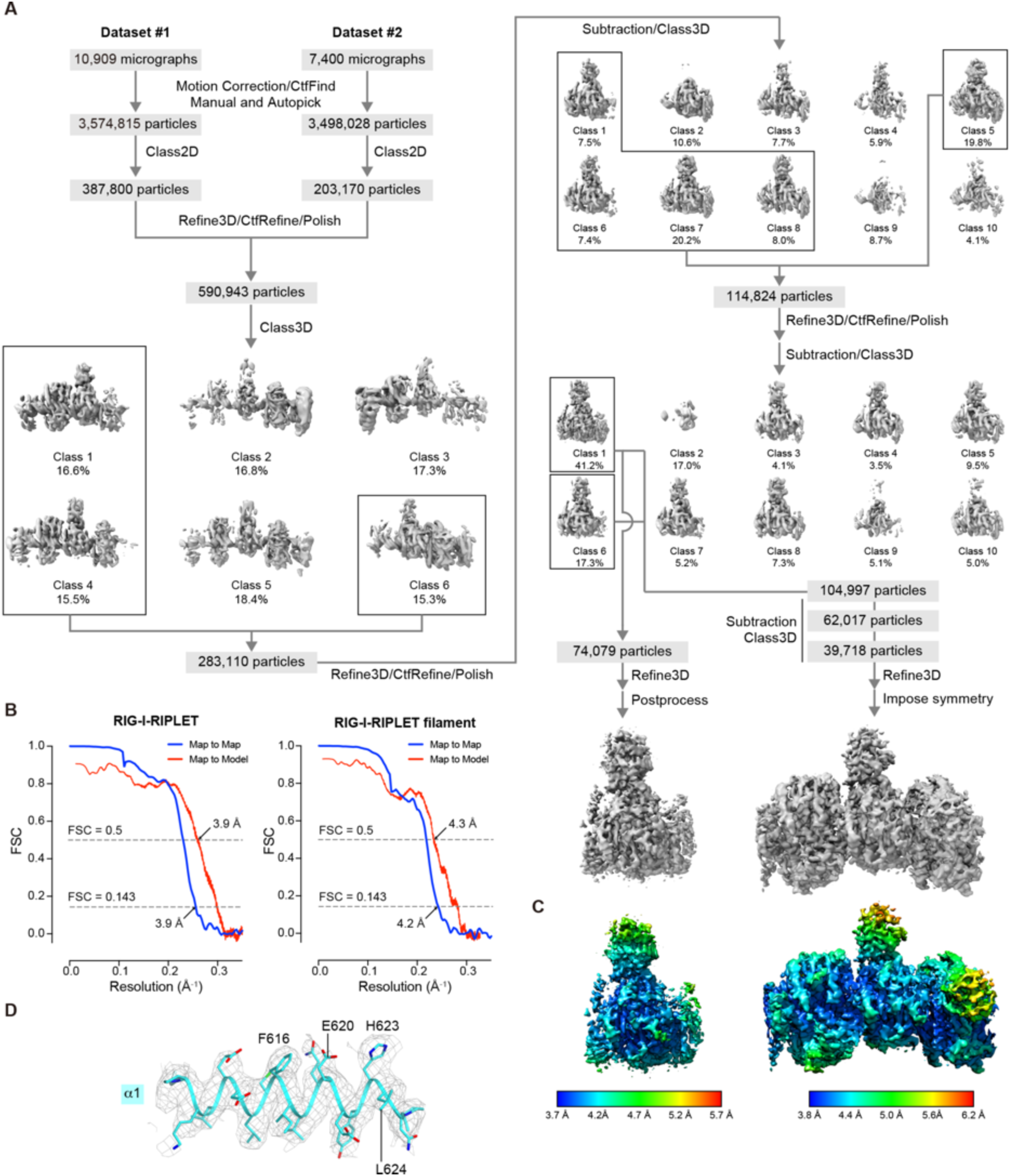
Cryo-EM data processing for the RIPLET^PSpry^:RIG-IΔN complex. (A) Cryo-EM image processing workflow (see details in Supplementary Methods). (B) Fourier shell correlation (FSC) curve for the monomeric (left panel) and filamentous (right panel) RIG-I:RIPLET complex. Map-to-Map FSC curve was calculated between the two independently refined half-maps after masking (blue line), and the overall resolution was determined by gold standard FSC=0.143 criterion. Map-to-Model FSC was calculated between the refined atomic models and maps (red line). (C) Local resolution for the maps of the monomeric (left panel) and filamentous (right panel) RIG-I:RIPLET complex. Local resolution was calculated by Relion, and resolution range was indicated according to the color bar. (D) Cryo-EM density map for the a1 helix of RIG-I.

**Figure S5.**
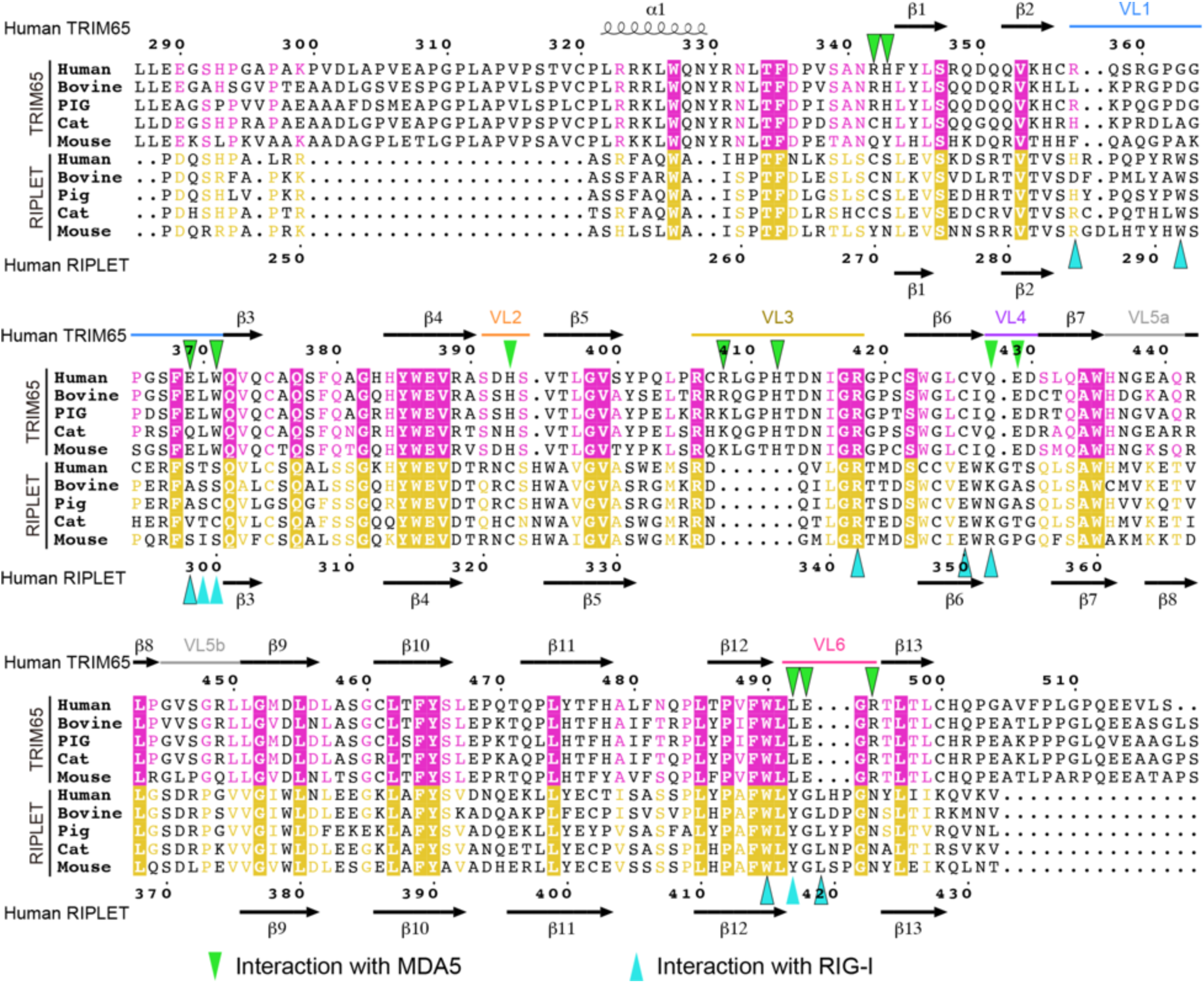
Sequence alignment of orthologs of TRIM65 and RIPLET near the MDA5/RIG-I interface. Residues involved in the interaction with MDA5 and RIG-I are indicated with triangles. Sequences were aligned using Clustal Omega (http://www.ebi.ac.uk/Tools/msa/clustalo), and alignment figures were generated using ESPript3 (http://espript.ibcp.fr/ESPript/ESPript).

**Figure S6.**
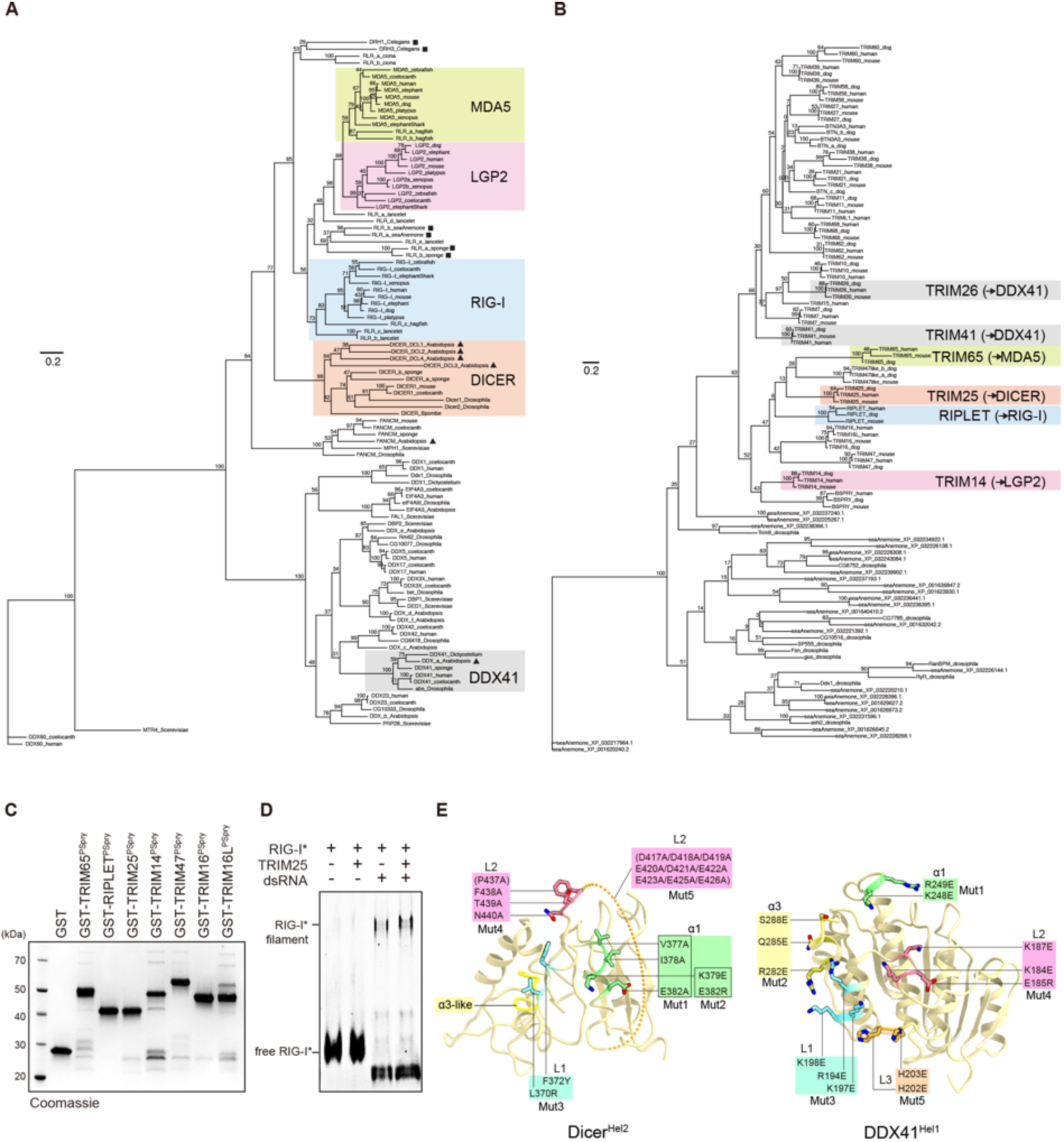
Analysis of related helicases and PSpry/Spry-containing proteins. (A) Phylogenetic trees of selected helicases and PSpry/Spry-containing proteins. The helicase domain phylogeny includes human representatives of the RLRs and their closest relatives DICER and FANCM, a number of DEAD-box helicases, and the outgroup DDX60, a Ski2-like helicase, for rooting. We added selected orthologs chosen to help put lower bound estimates on the age of the duplication events that gave rise to these diverse helicases. Because the plant *Arabidopsis thaliana* has clear orthologs (triangles) of human DDX41, FANCM and DICER, these genes must have diverged from one another in or before the common ancestor of all eukaryotes >1500 million years ago (mya) (Kumar et al., 2017). While we have not identified a plant ortholog for RIG-I/MDA5/LGP2, these proteins group with several *C. elegans, Nematostella vectensis* (sea anemone) and *Amphimedon queenslandica* (sponge) sequences (squares), indicating that they diverged from DICER >950mya (Zou et al., 2009). The scale bars represent 0.2 amino acid substitutions per site. Numbers at each node show bootstrap values (100 replicates). (B) The PSpry/Spry-domain phylogeny includes all TRIM proteins studied functionally in this manuscript and their close relatives, including human, dog and mouse orthologs where available. It is not intended to be a comprehensive TRIM phylogeny. We added all alignable Spry-containing proteins from *Drosophila melanogaster* and *Nematostella vectensis* (sea anemone) and display the tree with arbitrary rooting. Because the studied TRIM proteins group together to the exclusion of *Drosophila* and *Nematostella* sequences, we infer that the duplications giving rise to these TRIM proteins occurred <800mya. Interacting helicases are shown in parenthesis. The scale bars represent 0.2 amino acid substitutions per site. Numbers at each node show bootstrap values (100 replicates) (C) SDS-PAGE analysis of recombinant GST-PSpry proteins used in Figure 5A. (D) Native gel mobility shift assays to test RIG-I:TRIM25 interaction. Fluorescein-labeled (*) full-length RIG-I (600 nM) was incubated with TRIM25 (300 nM) in the presence and absence of 112 bp dsRNA (4 ng/μl). Fluorescein fluorescence was used for gel imaging. The result shows no binding between RIG-I and TRIM25, with or without dsRNA. (E) Mutations in Dicer and DDX41 used in Figure 6B-D. Mutated residues are mapped onto the structures of human Dicer Hel2 (PDB:5ZAL) and human DDX41 Hel1 (PDB:5GVR).

**Table S1.**
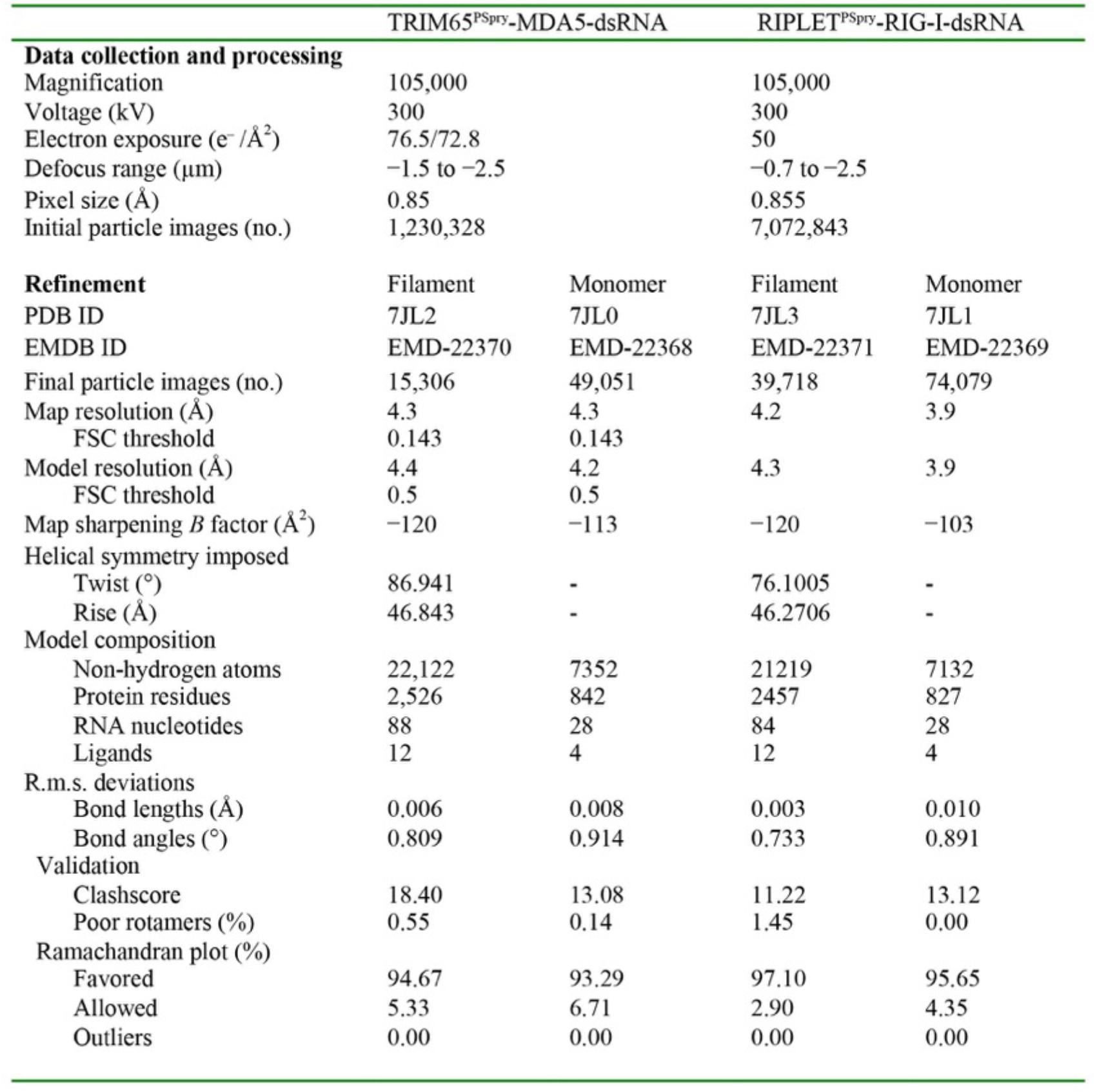
Cryo-EM data collection, refinement and validation statistics

**Table S2.**
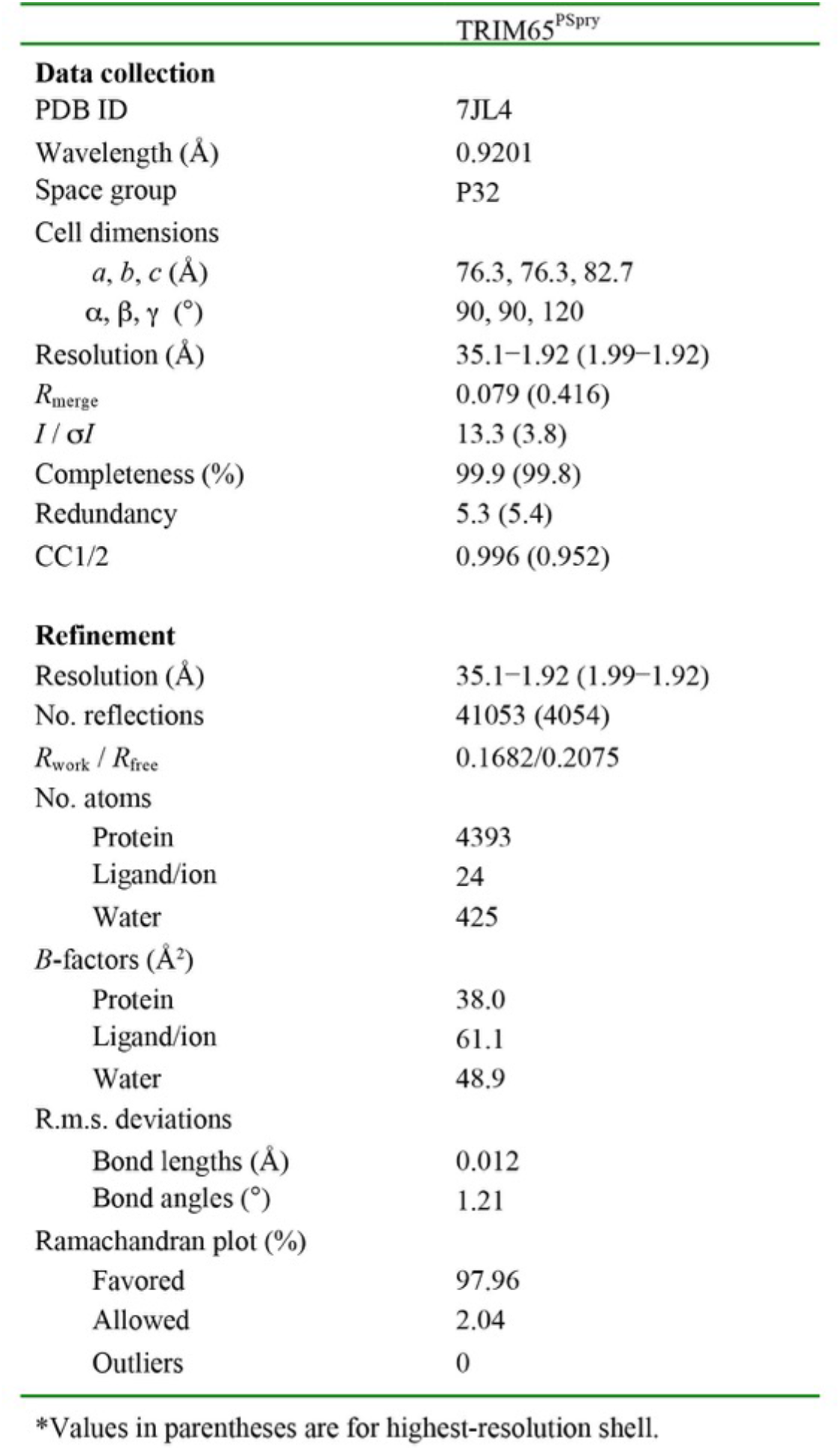
Crystallographic data and refinement statistics

## Material Preparation

### Plasmids

Mammalian expression plasmids for MDA5 and RIPLET were described previously (Cadena et al., 2019; Wu et al., 2013). Human *Trim65* was inserted in pFLAG-CMV4 vector between HindIII and ClaI sites for mammalian expression. Myc tagged Human *Rig-I* was inserted between KpnI and NotI sites of pcDNA3.1. Mammalian expression plasmid for MAVS has been described previously (Wu et al., 2014) and pcDNA4 expressing STING V155M variant was a kind gift from P. Kranzusch, Harvard Medical School. Baculovirus expression construct for LGP2 and TRIM65^CC-PSpry^ (residues 125–517) were cloned into the modified pFastBac1vector (Invitrogen), in which His_6_-Strep-tag and HRV 3C protease recognition site were inserted between the polyhedrin promoter and BamHI site. For bacterial expression of MDA5, the gene encoding MDA5, MDA5ΔN (residues 287–1025), helicase (residues 287–837) and α1 (residues 689–718) were inserted into pE-SUMO vector (LifeSensors) or the modified pGEX-6P-1 vector (GE healthcare) with N-terminal His6-tag. Bacterial expression plasmid for RIG-I was described previously (Cadena et al., 2019). The gene encoding RIG-IΔN (residues 204-925) was inserted into the pE-SUMO vector. For the plasmids of TRIM65^PSpry^-MDA5ΔN, MDA5ΔN-TRIM65^PSpry^ and RIPLET^PSpry^-RIG-IΔN, the two DNA fragments of PSpry (TRIM65 or RIPLET) and RLH (MDA5ΔN or RIG-IΔN) were cloned into pE-SUMO vector using InFusion HD (Clontech) together with a gene encoding a flexible linker comprised of 38 residues. All Hel2i2 constructs for MDA5 (residues 532–837), LGP2 (211–487) and Dicer (243–610) and GST-PSpry constructs for TRIM65 (295–517), RIPLET (249–442), TRIM14 (238–442), TRIM16 (346–564), and TRIM47 (401–638) were cloned into the modified pGEX-6P-1 vector with N-terminal His_6_-tag. All DDX41 constructs, Hel12 (155–578), Hel1 (155–401) and Hel2 (402–578) were cloned into the modified pGEX-6P-1 vector with N-terminal His_6_-tag.

TRIM14^CC-PSpry^ (residues 59–442) was cloned into the pE-SUMO vector. All bacterial expression constructs for TRIM25, TRIM26 and TRIM41 were cloned into the modified pE-SUMO vector, in which His_6_-Strep-tag and HRV 3C protease recognition site were inserted after the region encoding SUMO. For expression construct of TRIM65^PSpry^ used for crystallization, the gene encoding the residues 312-504 of TRIM65 were cloned into the modified pMal-c2x vector (NEB), in which N-terminal His_6_-tag and the HRV 3C protease recognition site were inserted into 5’-upstream a maltose binding protein sequence and a multiple cloning site, respectively. All mutations were introduced by a PCR-based method, and the sequences were confirmed by DNA sequencing.

### RNAs

Double-stranded RNAs (dsRNAs) used in this study were prepared by *in vitro* T7 transcription as described previously (Peisley et al., 2011). The templates for RNA synthesis were generated by PCR amplification. The sequences of 42, 112 and 1012 bp dsRNAs were taken from the first 30, 100 and 1000 bp of the MDA5 gene, respectively, flanked by 5’-gggaga and 5’-tctccc. The sequence for 15 bp dsRNAs was 5’-agggcccgggaugcu. The two complementary RNA strands were co-transcribed, and the duplex was purified using 8.5% acrylamide (112 bp) or 6% acrylamide (1012 bp) gel electrophoresis. RNA was gel-extracted using the Elutrap electroelution kit (Whatman), ethanol precipitated, and stored in 20 mM HEPES pH 7.0. For 112 bp dsRNA with a 5-nt 5’ overhang (5’ovg 112 bp dsRNA), 5-nt sequence (GGGTT) was inserted between T7 promoter sequence and the MDA5 gene of the template DNA transcribed for the negative strand RNA, and the two complementary RNA strands were transcribed and purified separately. The purified negative strand RNA was treated with Calf intestinal alkaline phosphatase (CIP, NEB) to remove 5’-triphosphate moiety, and was re-purified using QIAquick Nucleotide removal kit (QIAGEN). The dephosphorylated RNA was annealed with the 112-nt positive strand RNA in 50 mM NaCl and 10 mM EDTA. Qualities of RNAs were analyzed by 1X TBE polyacrylamide gel and the RNA duplex integrity was established using Acridine Orange staining (Sigma Aldrich) (Mu et al., 2018). For 3’-labeling with Cy5, 112 bp dsRNA was oxidized with 0.1 M sodium meta-periodate (Pierce) overnight in 0.1 M NaOAc pH 5.4. The reaction was quenched with 250 mM KCl, buffer exchanged using Zaba desalting columns (Thermo Fischer) into 0.1 M NaOAc pH 5.4 and further incubated with 0.1 M Cy5-Monohydrazide (GE Healthcare) for 4–6 hr at RT.

### Cell lines

The TRIM65-KD cells were generated using the method described previously (Chavez et al., 2015). The TRIM65-specific guide RNA sequences (<u>Forward</u>: TGCCGAGCGCCTCAAGCGCG; <u>Reverse</u>: CGCGCTTGAGGCGCTCGGCA) were inserted in pSB700 vector (Addgene) using BsmBI site. The plasmid was then transfected in HEK293T cells in a 6-well plate along with psPAX2 (Addgene) and pMD2.G (Addgene) packaging vectors in a ratio of 5:2.5:1. The lentiviral particles were harvested 48 h post-transfection and filtered using a 0.45 μ filter. The virus was then transduced into Cas9-stable HEK293T cells (generous gift from G. Church, Harvard Medical School) and allowed to grow in the presence of 3 μg/ml puromycin, 2.5 μg/ml blasticidin and 10 μg/ml hygromycin. For parental WT cells, virus derived from pSB700 empty vector was used and the rest of the procedure was the same as that for knockdown cells. 7 days after transduction, monoclonal cell masses were picked and screened for knockdown of TRIM65 using Western blot analysis. RIG-I^-/-^, RIPLET^-/-^ and their respective parental WT HEK293T cell lines were obtained from the Hou lab (Shi et al., 2017).

## Protein expression and purification

TRIM65 protein was expressed in HEK293T cells using polyethylenimine (PEI, Polysciences). The harvested cells were lysed by Dounce homogenizer in hypotonic buffer (10 mM KCl, 10 mM Tris pH7.5 and 1.5 mM MgCl_2_) supplemented with 1 mM PMSF and mammalian protease inhibitor cocktail (G-Biosciences). The lysed cells were centrifuged, and the supernatant was applied to anti-FLAG M2 agarose beads (Sigma-Aldrich). The protein was eluted with 3X FLAG peptide (Sigma-Aldrich), and buffer was exchanged with 50 mM HEPES pH7.5, 150 mM NaCl and 2 mM DTT using Zeba desalting columns, 40 kDa molecular-weight cutoff (Thermo Scientific) to remove the 3X FLAG peptide. LGP2 protein was expressed in Sf9 cells using the Bac-to-Bac baculovirus expression systems (Invitrogen). Sf9 cells were infected in HyClone SFX Insect cell culture media (GE Healthcare) for 48 h at 27°C. The harvested cells were lysed by sonication and centrifuged. The supernatant was applied to Ni-NTA agarose (QIAGEN) and the eluted protein was treated with HRV 3C protease at 4°C overnight to cleave the N-terminal tag. The protein was further purified by chromatography on HiTrap Heparin (GE Healthcare) and Superdex 200 Increase 10/300 (GE Healthcare) columns. For TRIM65^CC-PSpry^ (residues 125–517), expression and lysis were done in the same manner as LGP2, while protein purification was done by chromatography on Strep-Tactin Superflow (IBA). The eluted protein was treated with HRV 3C protease, and further purified by size-exclusion chromatography on Superdex 200 Increase 10/300 columns. All other proteins were expressed in *Escherichia coli* Rosetta2 (DE3) (Novagen) by inducing with 0.25 mM isopropyl β-D-thiogalactopyranoside (IPTG) at 18°C overnight. Cells were lysed by high-pressure homogenization using an Emulsiflex C3 (Avestin). The N-terminally His_6_-SUMO-tagged MDA5, RIG-I, and its variants were purified by chromatography on Ni-NTA agarose. The eluted proteins were incubated with Ulp1 protease at 4°C for 6 h to cleave the His_6_-SUMO-tag. The proteins were further purified by chromatography on HiTrap Heparin and Superdex 200 Increase 10/300 columns. All His_6_-GST-tagged proteins were purified by a combination of chromatography on Ni-NTA agarose and HiTrap Heparin/HiTrap Q (GE Healthcare) columns, followed by size-exclusion chromatography on Superdex 200 Increase 10/300. The CC-PSpry domains for TRIM14 (residues 59–442), TRIM25 (residues 189–630), TRIM26 (residues 134–539) and TRIM41 (residues 258–630) were expressed as a fusion protein with N-terminal His_6_-SUMO-tag, and purified by a combination of chromatography on Ni-NTA agarose, Resource Q/Resource S (GE Healthcare) and Superdex 200 Increase 10/300 columns. The CC domains (TRIM25: residues 189–362, TRIM26: residues 134–270 and TRIM41: residues 258–391) and the PSpry domains (TRIM25: residues 447–630, TRIM26: residues 271–539 and TRIM41: residues 392–630) were purified by a combination of chromatography on Ni-NTA agarose and Superdex 200 Increase 10/300 columns. TRIM65^PSpry^ (residues 295–517) and its crystallization construct (312–504, C356L, C407S and C421S) were expressed as a fusion protein with N-terminal His6-maltose binding protein (MBP)-tag. The proteins were purified by Ni-NTA agarose and treated with HRV 3C protease to cleave the His_6_-MBP-tag. The cleaved proteins were further purified by Resource S and Superdex 75 Increase 10/300 (GE Healthcare) columns. For N-terminal fluorescent labeling of MDA5, TRIM65 or LGP2 and their variants, the proteins (0.5–2 mg/ml) were incubated with 100 μM peptide (LPETGG) conjugated with fluorescein (Anaspec) and 1 mg/ml *S. aureus* sortase A (a gift from H. Ploegh, MIT) (Antos et al., 2009) at RT for 4 h away from light, followed by size-exclusion chromatography on Superdex 200 Increase 10/300 to remove sortase A. All purified proteins were frozen in liquid nitrogen and stored at –80°C until further use.

### Antibodies

The antibodies used for immunoblots were: Anti-FLAG-HRP (SIGMA A8592), Anti-HA (Cell Signaling 3724S), Anti-myc (Cell Signaling 2278S), Anti-TRIM65 (Invitrogen PA5-54459), Anti-RIG-I (Enzo ALX-210-932-C100), Anti-STING (Cell Signaling 13647S), Anti-Actin (Cell Signaling 8457L), Anti-MDA5 (Enzo ALX-210-935-C100), Anti-TRIM14 (Abcam ab185349, LS Bio C110434), Anti-TRIM25 (BD Biosciences 610570), Anti-TRIM26 (Santa Cruz sc-393832), Anti-TRIM41 (Abcam ab111580), Anti-Strep-tag (Invitrogen MA5-17283), Anti-mouse-IgG-HRP (GE Healthcare NA931V) and Anti-rabbit-IgG-HRP (Cell Signaling 7074P2).

## RT-qPCR

HEK293T cells were grown in 48-well plates in Dulbecco’s modified Eagle medium (Cellgro) supplemented with 10% fetal calf serum. The cells were transfected with indicated amounts of plasmid (MDA5 WT = 5 ng; MDA5 G495R = 10 ng; TRIM65 = 5–50 ng; RIG-I WT = 10 ng; RIPLET = 10 ng; MAVS = 25 ng; STING V155M = 750 ng) at around 70–80% confluency using Lipofectamine 2000 (Thermo Fisher) following manufacturer’s protocol. The plasmid dose for all point mutants were optimized using Western blot analysis to match the wild type expression levels. Similarly, for experiments requiring comparison of WT and KD cells, the plasmid dose was optimized to get similar expression levels across cell-lines. For experiments requiring RNA stimulation, the media was changed 6 h after the first transfection and the cells were then additionally transfected with 500 ng polyI:C (Invivogen) or 200 ng 42 bp dsRNA. The cells were lysed ~20 h after stimulation and the whole cell RNA was extracted with TRIzol reagent (Thermo Fisher) using Direct-zol RNA Miniprep kit (Zymoresearch) followed by cDNA synthesis using High Capacity cDNA reverse transcription kit (Applied Biosystems) according to the manufacture’s instruction. The cDNA was subjected to real-time PCR using a set of gene specific primers and SYBR Green Master Mix (Applied Biosystems) in the StepOne Real-Time PCR Systems (Applied Biosystems). The interferon signaling activity was quantified by measuring the increase in levels of IFNβ mRNA relative to the change in levels of 18S rRNA. Average values and standard deviations were calculated using Microsoft excel. Each datapoint was derived from 3–4 independent experiments and the *P* values were calculated using the twotailed unpaired Student’s t test in Microsoft excel. The *P* values are denoted by ** (*P* < 0.005) and ***(*P* < 0.001).

## Ubiquitination assay

Ubiquitination assay was performed as described previously (Cadena et al., 2019). Briefly, purified MDA5 or RIG-I (0.5 μM) was first incubated with 1012 bp or 112 bp dsRNA (2 ng/μl) respectively, in 20 mM HEPES pH 7.5, 50 mM NaCl, 1.5 mM MgCl_2_, 5 mM ATP and 2 mM DTT at RT for 30 min. The protein-RNA complex was then further mixed with 20 μM ubiquitin, 1 μM mE1, 5 μM Ubc13, 2.5 μM Uev1A and 0.25 μM TRIM65, and the mixture was incubated at 37°C for 30 min. The reaction was quenched with SDS loading buffer and analyzed on SDS-PAGE followed by anti-MDA5 or anti-RIG-I western blot.

## Native gel-shift assay

For experiments requiring dsRNA, 112 bp dsRNA was used unless mentioned otherwise. For detection purpose, either Cy5-labeled dsRNA or fluorescein-labeled protein was used (as mentioned in the relevant experiments). For experiments where Cy5-labeled dsRNA was used, MDA5 (or other RLHs) and/or their variants at 0.25 μM were first incubated with dsRNA (1 ng/μl) for 30 min in 20 mM HEPES pH 7.5, 100 mM NaCl, 1.5 mM MgCl_2_ and 2 mM DTT at RT with or without 2 mM ATP. This was followed by the addition of the TRIM proteins (TRIM65, RIPLET, TRIM14, TRIM16, TRIM25 or TRIM47) and/or their variants (as mentioned in the experiment). The mixture was further incubated at RT for 10 min and the samples were then analyzed on Bis-Tris native PAGE (Life Technologies). For experiments where fluorescin-labeled TRIM65 CC-PSpry was used, GST-tagged MDA5 variants (1.6 μM) were directly incubated with fluorescin-labeled TRIM65 CC-PSpry (0.8 μM) for 15 min at RT before analyzing on native PAGE. For experiments where fluorescin-labeled MDA5 was used, fluorescin-labeled MDA5 or its variants (0.6 μM) was incubated with TRIM65 (0.3 μM) in the presence or absence of 112 bp dsRNA (8 ng/μl) for 30 min, and the mixture was analyzed on native PAGE. The gels were imaged using either Cy5 or fluorescin fluorescence in iBright FL1000 (Invitrogen).

## Multi-Angle Light Scattering (MALS)

The molecular masses of TRIM65^CC-PSpry^ and TRIM65^PSpry^ were determined by MALS using a Superdex 200 Increase 10/300 column (GE Healthcare) attached to MiniDAWN detector (Wyatt Technology) in Phosphate Buffered Saline (PBS, Corning) buffer containing 2 mM DTT and the data were analyzed using ASTRA7.3.1 software (Wyatt Technology).

## Pull-down assay

GST-tagged helicase proteins (200 nM) and TRIM proteins (200 nM) were incubated in PBS buffer containing 2 mM DTT at 4°C for 2 h, followed by the addition of Glutathione Magnetic Beads (Pierce). The samples were further incubated at 4°C for 2 h with gentle rotation, and the beads were washed with PBS containing 2 mM DTT three times. The bound proteins were eluted by adding the buffer containing 50 mM Tris-HCl pH 7.5, 300 mM NaCl, 2 mM DTT and 50 mM glutathione (Sigma-Aldrich). The eluted samples were boiled with SDS-loading buffer at 96°C for 3 min and analyzed by immunoblotting using anti-Strep-tag or anti-TRIM14.

## Crystallization, X-ray data collection, processing and refinement

The purified human TRIM65^PSpry^ (residues 312–504, C356L/C407S/C421S) protein was used for crystallization. Three solvent exposed cysteine residues (Cys356/Cys407/Cys421) were mutated to enhance crystallization. To further facilitate crystallization, the lysine residues of the protein were methylated, as previously described (Walter et al., 2006). In brief, the protein was incubated with the dimethylamine-borane (ABC) complex and formaldehyde in 50 mM HEPES pH 7.5, 150 mM NaCl at 4°C overnight. The methylated TRIM65^PSpry^ protein was further purified by gel filtration chromatography on a Superdex 75 Increase column (GE Healthcare) and concentrated to 5 mg/ml using an Amicon Ultra-4 filter (10 kDa molecular-weight cutoff; Millipore). The methylated TRIM65^PSpry^ was crystallized at 18°C by the hanging drop vapor diffusion method. Crystals were obtained by mixing 1 μl of protein solution (5 mg/ml TRIM65^PSpry^, 10 mM Tris-HCl pH 8.0, 150 mM NaCl and 3 mM 2-mercaptoethanol) and 1 μl reservoir solution (0.1 M Bicine, pH 9.0 and 20% PEG3,350). Crystals were cryoprotected in reservoir solution supplemented with 25% glycerol, and were flash-cooled in liquid nitrogen. The X-ray diffraction data was collected on 17-ID beamline at the NSLSII and processed using XDS (Kabsch, 2010). The structure was solved by molecular replacement with Phaser (McCoy et al., 2007). The search model was built by the Phyre 2 web server (Kelley et al., 2015), using the PSpry domain of TRIM25 (PDB ID 6FLN) as a template. The model was automatically built using Autobuild (Terwilliger et al., 2008), followed by manual model building using COOT (Emsley et al., 2010) and structural refinement using PHENIX (Liebschner et al., 2019). The structure was validated using MolProbity (Williams et al., 2018). Data collection and refinement statistics are summarized in Extended Data Table 1.

## CryoEM sample preparation and data collection

TRIM65^PSpry^-MDA5ΔN (8.6 μM) was incubated with 1012 bp dsRNA (50 ng/μl) in 20 mM HEPES pH 7.5, 150 mM NaCl, 1.5 mM MgCl2, 2 mM DTT and 2 mM ADP:AlF_x_ (ADP, AlCl3 and NaF in a molar ratio of 1:1:3) at 4°C for 2 h to form the protein-RNA filament. The filament sample was diluted 2-fold and 3.5 μl of the diluted sample was applied to a freshly glow-discharged C-flat 400 mesh copper grid (CF-1.2/1.3, Electron Microscopy Sciences) at 4°C at 100% humidity and plunged into liquid ethane after blotting for 3 s using a Vitrobot Mark IV (FEI). The grids were screened at Pacific Northwest Centre for Cryo-EM, OHSU using Talos Arctica microscope (FEI). The grids that showed a good sample distribution and ice thickness were subjected to data collection on a Titan Krios G3i microscope (Thermo Fisher 20 Scientific) operated at 300 kV and equipped with K3 summit direct electron detector (Gatan) at the Harvard Cryo-Electron Microscopy Center for Structural Biology (Harvard Medical School). A total of 12,248 micrographs were recorded from 2 independent sessions in counting mode at a magnification of 105,000x and magnified pixel size 0.85 Å using SerialEM software (Mastronarde, 2005). Each movie comprised of 50 frames at a dose rate of 35.604 e^−^/Å^2^/s and an exposure time of 2.1 s resulting in a total dose of 74.8 e^−^/Å^2^. The data was collected in a desired defocus range of –1.5 to –2.5 μm.

RIPLET^PSpry^-RIG-IΔN (9.6 μM) was incubated with 112 bp dsRNA (48 ng/μl) at 37°C for 5 min in 20 mM HEPES pH 7.5, 150 mM NaCl, 5 mM MgCl_2_, 2 mM DTT and 1 mM ATP to form the protein-RNA filament. The reaction was quenched with 5 mM ADP:AlFx and applied to the EM-grid without dilution and blotted for 2 s. The data for RIPLET^PSpry^-RIG-IΔN was collected at National Cancer Institute’s National Cryo-EM Facility at the Frederick National Laboratory for Cancer Research using Titan Krios G3i microscope operated at 300 kV equipped with K3 summit direct electron detector (Gatan) in counting mode at 105,000x magnification and 0.855 Å pixel size. A total of 18,309 micrographs were recorded from 2 independent sessions with each movie comprising of 40 frames at a dose rate of 19.424 e^−^/ Å^2^ per s and an exposure time of 2.58 s, resulting in a total dose of 50.0 e^−^/ Å^2^. The data was collected in a desired defocus range of –0.7 to –1.4 μm.

## CryoEM data processing and structure refinement

All image processing was performed in RELION3.08 or RELION3.1 (Zivanov et al., 2018). The dose-fractionated movies were motion-corrected using MotionCor2 (Zheng et al., 2017). The contrast transfer function (CTF) was estimated with CTFFIND 4.1 (Rohou and Grigorieff, 2015). Particles were picked using the Autopick function in RELION (Zivanov et al., 2018). For TRIM65^PSpry^-MDA5ΔN, helical segments were extracted in a box size equal to 300 pixel with an inter-box distance of 46 Å. These particles were subject to several rounds of 2D classifications and selection in RELION 3.1 to get rid of badly aligned particles. The selected class averages were used for 3D refinement with helical reconstruction using a featureless cylinder as the starting reference map. The resulting 3D map included three monomers of TRIM65^PSpry^-MDA5ΔN helically arranged on the central dsRNA duplex. The resulting 3D model and particle set were subjected to per-particle defocus refinement, beam-tilt refinement, Bayesian polishing (Zivanov et al., 2019) and 3D classification. The selected class containing 143,063 particles were subjected to another round of per-particle defocus refinement and Bayesian polishing. To deal with structural heterogeneity of TRIM65^PSpry^, no-align 3D classification was performed using a mask covering the central monomer of TRIM65^PSpry^-MDA5ΔN-dsRNA duplex. The best class contained 49,051 particles and the resulting 3D map was subjected to 3D refinement and postprocessing, yielding a map with a global resolution of 4.3 Å, according to the Fourier shell correlation (FSC) = 0.143 criterion (Rosenthal and Henderson, 2003). For filamentous TRIM65^PSpry^-MDA5ΔN, no-align 3D classifications were performed for all three monomers of TRIM65^PSpry^-MDA5ΔN-dsRNA duplex. The resulting 15,306 particles yielded a map with a global resolution of 4.3 Å after 3D refinement with helical reconstruction followed by imposing helical symmetry on the unfiltered half-maps.

For RIPLET^PSpry^-RIG-IΔN, helical segments were extracted in a box size equal to 296 pixel with an inter-box distance of 46 Å. After several rounds of 2D classifications, the particles were subjected to 3D refinement, per-particle defocus refinement, beam-tilt refinement, Bayesian polishing, and 3D classification. The selected classes containing 283,110 particles were further classified by no-align 3D classification using a mask covering the central monomer of RIPLET^PSpry^-RIG-IΔN-dsRNA duplex, yielding 114,824 particles. These particles were subjected to 3D refinement, per-particle defocus refinement, beam-tilt refinement, Bayesian polishing, and no-align 3D classification. The best class contained 74,079 particles and the resulting 3D model was subjected to 3D refinement and postprocessing, yielding a map with a global resolution of 3.9 Å. For filamentous RIPLET^PSpry^-RIG-IΔN, no-align 3D classification was performed for all three monomers of RIPLET^PSpry^-RIG-IΔN-dsRNA duplex. The resulting 39,718 particles yielded a map with a global resolution of 4.2 Å after 3D refinement with helical reconstruction followed by imposing helical symmetry on the unfiltered half-maps. The local resolution was estimated by RELION. The data processing scheme is summarized in Extended Data Fig.2 and 3.

For TRIM65^PSpry^-MDA5ΔN-dsRNA duplex, the crystal structure of MDA5 complexed with 12 bp dsRNA (PDB ID: 4GL2) and TRIM65^PSpry^ were used as starting atomic model. These models were docked as a rigid body into the postprocessed EM density map in UCSF Chimera (Pettersen et al., 2004), and built manually against the density map using COOT (Emsley et al., 2010). The model was refined using phenix.real_space_refine (Liebschner et al., 2019), with the restrains for side chain rotamer, Ramachandran, secondary structure and base pair/stacking. For the filamentous TRIM65PSpry-MDA5ΔN-dsRNA duplex, the non-crystallographic symmetry (NCS) restraints were applied between each monomer of TRIM65^PSpry^-MDA5ΔN in real space refinement. To generate the filamentous trimer model, 3 repeats of the 14 bp dsRNA were used since there was no information about the RNA sequence in the maps due to helical symmetry averaging.

For RIPLET^PSpry^-RIG-IΔN-dsRNA duplex, the initial model of RIPLET^PSpry^ was built by the Phyre 2 server (Kelley et al., 2015), using the PSpry domain of TRIM25 (PDB ID: 6FLN) as a template. The modeled RIPLET^PSpry^ and the crystal structure of RIG-I complexed with 14 bp dsRNA (PDB ID: 5E3H) were docked into the postprocessed EM density map. The model building and structural refinement were performed as for TRIM65^PSpry^-MDA5ΔN-dsRNA duplex. The structure validation was performed using MolProbity (Williams et al., 2018) from the PHENIX package. The curve representing model vs. full map was calculated, based on the final model and the full map. The statistics of the 3D reconstruction and model refinement are summarized in Extended Data Table 2. All molecular graphics figures were prepared with CueMol (http://www.cuemol.org) and UCSF Chimera (Pettersen et al., 2004).

## Negative-stain electron microscopy

The samples of filamentous TRIM65^Spry^-MDA5ΔN and RIPLET^PSpry^-RIG-IΔN were prepared as described above. The samples were diluted 10-fold with 20 mM HEPES pH 7.5, 150 mM NaCl, 1.5 mM MgCl_2_ and 2 mM DTT, and was then immediately adsorbed to freshly glow-discharged carbon-coated grids (Ted Pella) and stained with 0.75% uranyl formate as described (Ohi et al., 2004). Images were collected using a JEM-1400 transmission electron microscope (JEOL) at 50,000x magnification.

## Phylogenetic analysis

Sequence collection and phylogenetic analysis were performed iteratively, using phylogenies to select additional species to add to the analysis, and/or additional query sequences to diversify the range of blast searches. To identify sequences, we used a combination of HMMER (v.3.1b2, http://hmmer.org/) and BLASTP (Altschul et al., 1997) searches. BLASTP searches used diverse queries across the trees shown, against either local databases representing the downloaded proteomes of selected species, or against NCBI’s non-redundant (NR) database, using the ‘entrez_query’ option to focus output on species groups of interest. HMMER searches used the hmmsearch algorithm and PFAM’s Hidden Markov Models (HMMs) for the SPRY (PF00622) and PRY (PF13765) domains, as well as HMMs we generated using hmmbuild from more focused alignments of only the TRIM proteins of interest. The target databases for HMMER searches were downloaded proteomes of species of interest, obtained from NCBI, UCSC, Ensembl, FlyBase, SGD, and Wormbase.

For phylogenies, sequences were aligned using MAFFT (Katoh and Standley, 2013) with the ‘-- leavegappyregion’ option. Alignments were manually trimmed to retain only well-aligned regions, poorly aligning sequences were removed, and the prottest (Darriba et al., 2011) algorithm was used to select an amino acid substitution model, which in both cases was the WAG model. Alignments were then used to generate a maximum likelihood phylogeny with the phyml (Guindon et al., 2010) algorithm, using empirical amino acid frequencies, and 100 bootstrap replicates to estimate the confidence of each node.

## Notes

### Competing Interest Statement

The authors have declared no competing interest.

